# Threat reduction must be coupled with targeted recovery programmes to conserve global bird diversity

**DOI:** 10.1101/2025.01.09.632120

**Authors:** Kerry Stewart, Chris Venditti, Carlos P. Carmona, Joanna Baker, Chris Clements, Joseph A. Tobias, Manuela González-Suárez

## Abstract

Biodiversity is undergoing severe declines while threats to natural populations continue to escalate worldwide. Ambitious international commitments have been made to preserve biodiversity, with the goal of preventing extinctions and maintaining ecosystem resilience, yet the efficacy of large-scale protection remains unclear. Here we use a trait-based approach to show that global actions – such as the immediate abatement of all threats across at least half of species ranges for ∼10,000 bird species - will only prevent half of projected species extinctions and functional diversity loss attributable to current and future threats in the next 100 years. Nonetheless, targeted recovery programmes prioritizing protection of the 100 most functionally unique threatened birds could avoid 68% of projected functional diversity loss. Actions targeting ‘habitat loss and degradation’ will prevent the greatest number of species extinctions and proportion of functional diversity loss relative to other drivers of extinction, whereas control of ‘hunting and collection’ and ‘disturbance and accidental mortality’ would save fewer species but disproportionately boost functional richness. These findings show that conservation of avian diversity requires action partitioned across all drivers of decline, and highlights the importance of understanding and mitigating the ecological impacts of species extinctions predicted to occur even under optimistic levels of conservation action.

## Main

Biodiversity is declining at an unprecedented rate with implications for ecosystem functioning and delivery of ecosystem services^1,2^. Human activity has led to widespread decline in the extent and structural condition of ecosystems, and changes in community trait composition^3^. High functional diversity – the diversity of traits which describe an organism’s ecological niche – has been associated with greater ecosystem functioning^4,5^, more reliable ecosystem service delivery^6^, and greater ecosystem resilience^7,8^. Therefore, changes in community composition could undermine the persistence of natural communities. Owing to the potential importance of functional diversity in supporting ecosystem function and resilience^9^, identifying effective measures for conserving functional diversity alongside species richness is paramount^10^.

Ambitious policies and substantial conservation resources have been dedicated to halting and reversing biodiversity loss by reducing the impact of threats^11^. However, it remains unclear whether functional diversity can be effectively preserved through a programme designed to alleviate threats at a large-scale – a strategy referred to as threat abatement^12^. Previous studies rarely go beyond isolated analyses of single threats and their impacts on species richness, or broader syntheses of the coverage of conservation targets (see ^13,14^). The main alternative to threat abatement strategies are direct management interventions such as breeding programmes and translocations. These measures can be effective^15,16^, particularly for rare species or those that are vulnerable to human pressures^17^. However targeted recovery programmes, including ex-situ conservation and in-situ measures to boost species survival and success, are often prohibitively expensive^18^ limiting their application as a global conservation strategy^17^. It is likely that conserving bird diversity can only be achieved with a combination of large-scale protection through threat abatement coupled with targeted species recovery programmes^15,19^, yet the extent to which abatement can reduce the need for intensive management to boost species population and reproductive success remains unclear.

Here, we use a trait-based approach to evaluate how much biodiversity and associated ecological function could be protected under different global conservation strategies. We assess the likely success of strategies focusing on the abatement of current and future drivers of extinction and estimate whether shortfalls in efficacy can be countered through targeted species recovery programmes. We used a phylogenetic generalized linear mixed model (PGLMM) to predict species extinction risk based on threats listed by the IUCN, accounting for non-independence geographically and across the avian tree of life. We quantify the importance of conserving unique species, which make a disproportionate contribution to the global diversity of form and function in birds.

## Projected species extinctions and functional diversity loss

We projected expected bird extinctions for the next 100 years based on IUCN Red List threat data^20^. We fitted a PGLMM implemented in a Bayesian Markov Chain Monte Carlo framework that predicted species assignment to Red List category with 86.8% accuracy (see Supplementary Information), using data on threat scope and severity, and including random effects describing the spatial and phylogenetic relationships among species. We then stochastically projected species extinctions based on expected probabilities of extinction for each Red List category (see Methods).

In the baseline extinction scenario, we assumed that human activity and natural threats will continue to impact bird populations as currently listed. In this scenario, we predict that 5.2 ± 0.2% (mean ± SD) of the 9873 extant birds studied will go extinct in the next 100 years (517 ± 19 species) (Fig. 1); more than three times the recorded number of bird extinctions since 1500. This figure falls within the range of previously predicted bird extinctions, ranging from Monroe et al.^21^ who predicted 226–589 species extinctions in the next 500 years, to Andermann et al.^22^ who predicted 669-738 species extinctions in the next 100 years. Extinctions on this scale are expected to fundamentally alter the global bird assemblage, potentially reducing functional diversity^23,24^.

**Figure 1.**
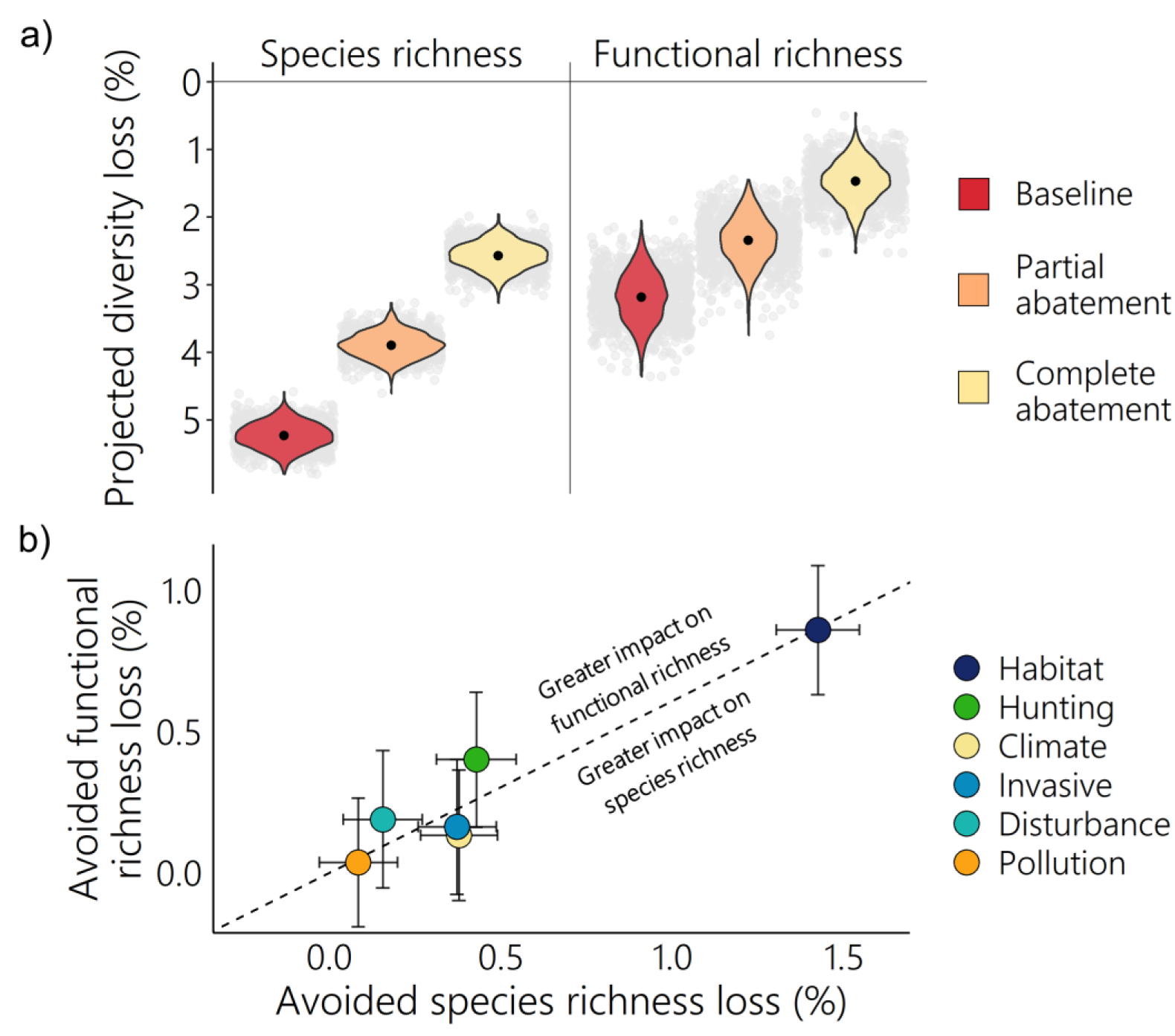
Projected loss of avian diversity in the next 100 years. **a,** Loss in species richness and in functional richness under three scenarios: baseline extinction, partial abatement of all drivers of extinction, and complete abatement of all drivers of extinction. Black points show mean loss across 1000 iterations for each scenario, with variation in those points show by their distribution (violin plots) and the individual values (grey dots). **b,** Diversity loss avoided under driver-specific complete abatement of six major drivers of extinction (circles represent the mean and error bars represent 0.5 standard deviations). Note that in some iterations loss avoided could be negative as more diversity was lost with driver-specific abatement than under the baseline scenario. The dotted diagonal line shows mean functional richness loss per species richness loss under complete abatement of all threats. Drivers above this line show greater avoidance of functional richness loss per species richness loss avoided, relative to the mean across all drivers of extinction. Hunting = hunting and collection; Climate= climate change and severe weather; Invasive = invasive species, genes and disease; Disturbance = disturbance and accidental mortality. Analyses based on 9873 species (of which 2087 species currently listed as Near Threatened or in threatened categories were modelled and could have reduced extinction risk in the abatement scenarios). 1000 iterations were run for each extinction scenario.

To quantify projected change in functional diversity in the world’s avifauna (*n* = 9873) we used published data on 11 continuous morphological traits that collectively capture bird ecological niches through their well-established association with diet, dispersal and habitat^25–27^. These traits were summarised using the first three axes produced by phylogenetic principal component analysis, which explained 87.2% of variance in the dataset, providing an overview of global avian functional diversity (Extended Data Fig. 1; see Methods). We estimate functional diversity using probabilistic hypervolumes^28^, which can be applied to multidimensional data and are less sensitive to extreme trait values than other methods such as convex hulls^29^. We quantify the volume of trait space occupied by the current global avian assemblage (*n*=9873), and under future extinction scenarios. Under the baseline extinction scenario, functional diversity was projected to decrease by 3.2 ± 0.4% in the next 100 years, relative to present-day functional diversity. This is likely a conservative estimate that only reflects loss in three-dimensional functional space and ignores internal erosion of the space^23,30^. Projected functional diversity loss varied between 2.4 ± 0.9% and 3.8 ± 0.4% when measured with two and four-dimensions respectively (see Supplementary information).

## Large-scale protection from the drivers of extinction

Threat abatement could prevent species extinctions and reduce functional diversity loss. However, it is unclear to what extent biodiversity loss can be avoided, and what scale of action is required to prevent species extinctions and functional diversity loss altogether. Using our PGLMM model we predicted how extinction risk would change under three management scenarios that reflect varying levels of threat abatement (see Methods, Extended Data Fig. 2). Complete abatement involved removal of all direct drivers of extinction across the entirety of all species ranges, partial abatement involved removal of all direct drivers of extinction across at least half of all species ranges (threat spatial scopes downgraded to “*Minority <50%”*), and minimal abatement involved removal of all direct drivers of extinction across at least 10% of all species ranges (threat spatial scopes downgraded to “*Majority 50-90%”*).

Under the complete abatement scenario, half of biodiversity loss predicted under the baseline scenario could be prevented (Fig. 1, Extended Data Table 1). However, an average loss of 2.6 ± 0.2% of species richness (254 ± 19 species) and 1.5 ± 0.3% of functional richness remained. Because our model did not include Least Concern species so could not predict Least Concern status, it could overestimate diversity loss. We show that this was not the case however, as we obtained similar estimates (241 ± 18 species extinctions; 1.4 ± 0.3% functional diversity loss) when the analyses were rerun assuming a lower extinction probability for the Near Threatened category to match that of the Least Concern category (1 x 10^-6^ extinctions per species per year; see Supplementary information).

Some extinctions were not preventable even with complete abatement, so were not attributable to current and future drivers of extinction. These extinctions could reflect particularly vulnerable species which have high extinction risk despite being affected by few threats, as well as species that were severely affected by past threats which can no longer be managed or abated. The model captures variation in species vulnerability to extinction that is not described by threats through spatial and phylogenetic random effects. With the expectation that not all the variables affecting extinction risk will be known or measured, but many are phylogenetically or spatially correlated (such as life history or habitat quality), random effects were used to capture variation in species extinction risk due to factors associated with where a species lives or their evolutionary history.

The Cebu Flowerpecker (*Dicaeum quadricolor*) is a Critically Endangered species predicted to be at risk of extinction even under complete abatement due to factors associated with where it lives. The Cebu Flowerpecker has a very small and declining population (60-70 individuals)^31^ and our model suggests that the species will most likely go extinct without complementary measures such as a habitat restoration or exsitu conservation. Another threatened species, the Spectacled Petrel (*Procellaria conspicillata*), is unlikely to benefit from large-scale protection because its main threat – fishing bycatch – has a negligible impact on its population^32^. It remains at risk of extinction owing to factors associated with its evolutionary history.

Our finding that even large-scale and ambitious actions leading to the removal of all present, future, and likely-to-return threats will fail to prevent almost half of projected species extinctions challenges some of the key assumptions of global metrics used to track conservation progress. For example, the Species Threat Abatement and Restoration (STAR) metric^33^ is based on IUCN Red List data on threat scope and severity but assumes that complete threat abatement will allow the vast majority of species to be downgraded to Least Concern, an assumption which our findings do not support. While Mair et al.^33^ acknowledge that some species may require restoration to be downgraded to Least Concern, our results suggest that many species will require conservation measures in addition to threat abatement. Even when species are affected by no current or future threats they may still be threatened with extinction, owing to ongoing population decline (particularly in populations which are no longer self-sustaining), high vulnerability to stochastic events due to small population or range size, or reduced fitness due to severe population decline in the past^34^.

Partial abatement was somewhat effective at reducing avian diversity loss, preventing 26 ± 4% of projected species extinctions and 26 ± 13% of projected functional diversity loss (Fig. 1, Extended Data Table 1). Species that were experiencing severe declines but were affected by few threats showed greatest reduction in extinction risk under partial abatement. The Green-faced parrot finch (*Erythrura viridifacies*) and the Saint Vincent parrot (*Amazona guildingii*) responded particularly well to partial abatement, with a reduction in extinction risk that was almost as great as the reduction in extinction risk under complete abatement. Of the diversity loss that was attributable to the drivers of extinction (diversity loss under complete abatement), approximately half was prevented through partial abatement (50 ± 7% of species extinctions and 49 ± 26% of functional diversity loss). However, a 3.9% decrease in species richness (385 ± 18 species) and 2.3 ± 0.3% decrease in functional richness remained. The minimal abatement scenario prevented only a small proportion of biodiversity loss (Extended Data Table 1). Using different traits to quantify functional diversity did not affect our conclusions (see Supplementary information) as we obtained similar results when we used phylogenetic principal components constructed from three-dimensional scans of beak morphology^35^, and when the first phylogenetic principal component (largely describing variation in body size) was removed.

Protecting species from the drivers of extinction does not provide a comprehensive solution to biodiversity loss in the near future without additional measures such as targeted species recovery programmes, habitat restoration, and prioritisation of protection in important areas^36^. This finding is consistent with previous studies assessing biodiversity impacts of future conservation and mitigation scenarios. Leclere et al^.37^ found that while it was possible to bend the curve of biodiversity loss with an integrated strategy, protected area management and expansion to avert the impact of habitat loss and degradation were insufficient to avoid more than 50% of projected biodiversity loss on average in biodiversity rich regions. Similarly, Pereira et al.^38^ predict that in a 2015-2050 scenario of strong land use and climate change mitigation globally, rates of biodiversity loss would decrease, but biodiversity would continue to decline. The abatement scenarios explored here represented significant management efforts with optimistic assumptions about their impact and uptake. We assumed that the drivers of extinction, and the species declines caused by these drivers, could be halted immediately, and that all drivers of extinction could be alleviated, including climate change, which arguably may be difficult to mitigate with site-based protection. Even in these ambitious and optimistic scenarios, we predicted that over half of projected species extinctions and loss of functional diversity in the next 100 years would occur anyway.

Projected loss of functional diversity was not evenly distributed across functional space (Fig. 2, Fig. 3). Areas of trait space with high pPC1 values (generally larger birds) were predicted to show the greatest proportional losses under the baseline extinction scenario. Complete abatement was predicted to reduce loss across functional space but was less effective in a region of high pPC1 and pPC2 values, predominantly occupied by large aquatic predators.

**Figure 2.**
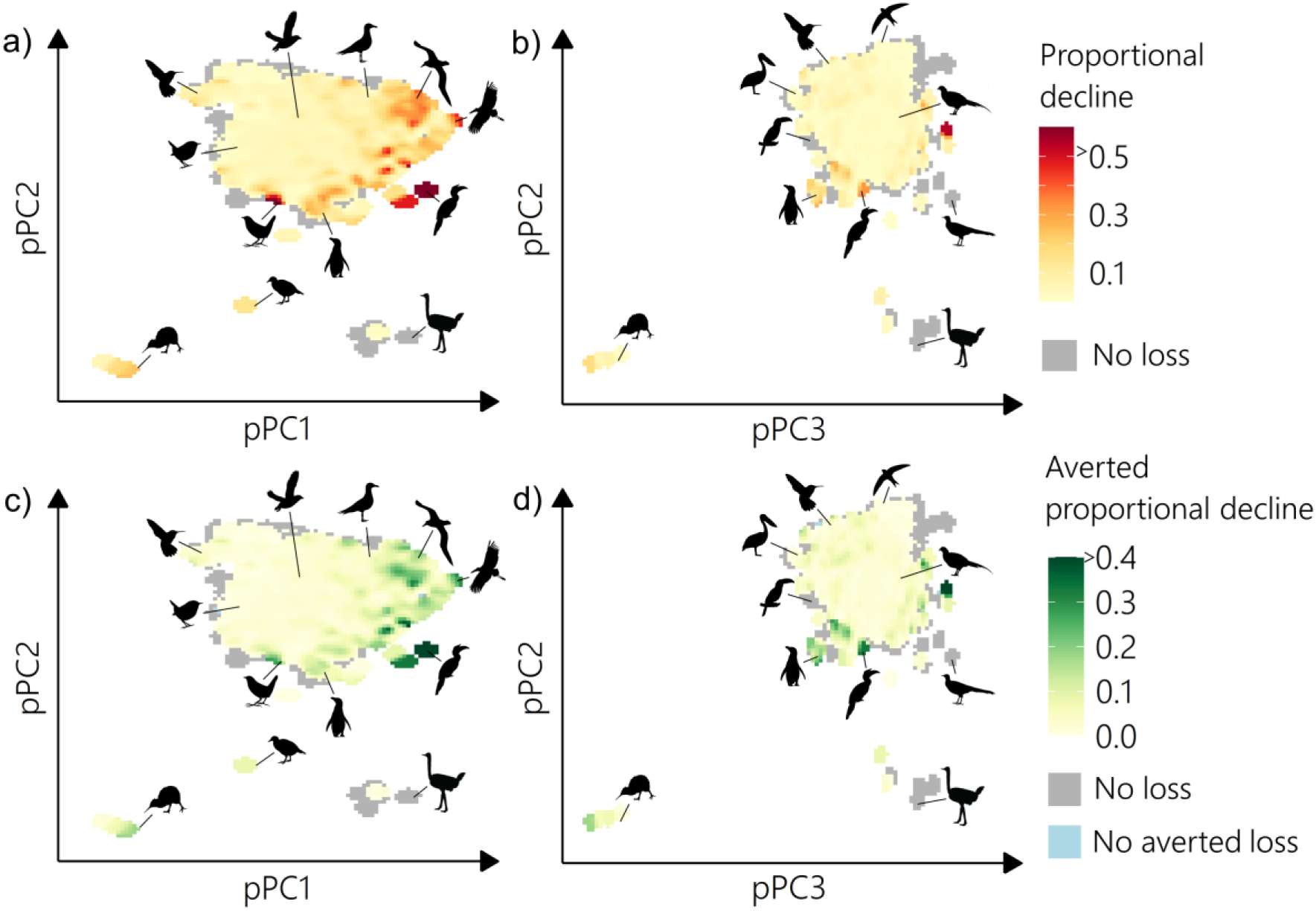
Change in occupation of morphospace under extinction and conservation. pPC1 is a descriptor of body size, explaining variance in wing length, beak length (nares and culmen), beak depth, beak width, body mass, tail length, tarsus length and Kipp’s distance. pPC2 is associated with wing morphology, explaining variance in hand wing index and Kipp’s distance. pPC3 explains variance in beak length and shape and tail length. a) and b) show predicted proportional decline in functional trait space occupation in the next 100 years under the baseline extinction scenario with respect to pPC1 and pPC2 (a) and pPC3 and pPC2 (b). Grey areas mark where there was no decrease in functional trait space occupation or there was a decrease in functional trait space occupation that was not preventable by managing drivers of extinction. Bottom panels (c and d) show averted proportional decline under the complete abatement scenario in which all current and future drivers of extinction, including those that occurred in the past and are likely to return, are removed entirely, for pPC1 and pPC2 (panel c) and for pPC2 and pPC3 (panel d). Grey shows areas where no functional diversity loss was projected, and blue shows areas where no functional diversity loss was avoided under complete abatement. Analyses based on 9873 species (of which 2087 species currently listed as Near Threatened or in threatened categories were modelled and could have reduced extinction risk in the abatement scenarios). 1000 iterations were run for each extinction scenario.

**Figure 3.**
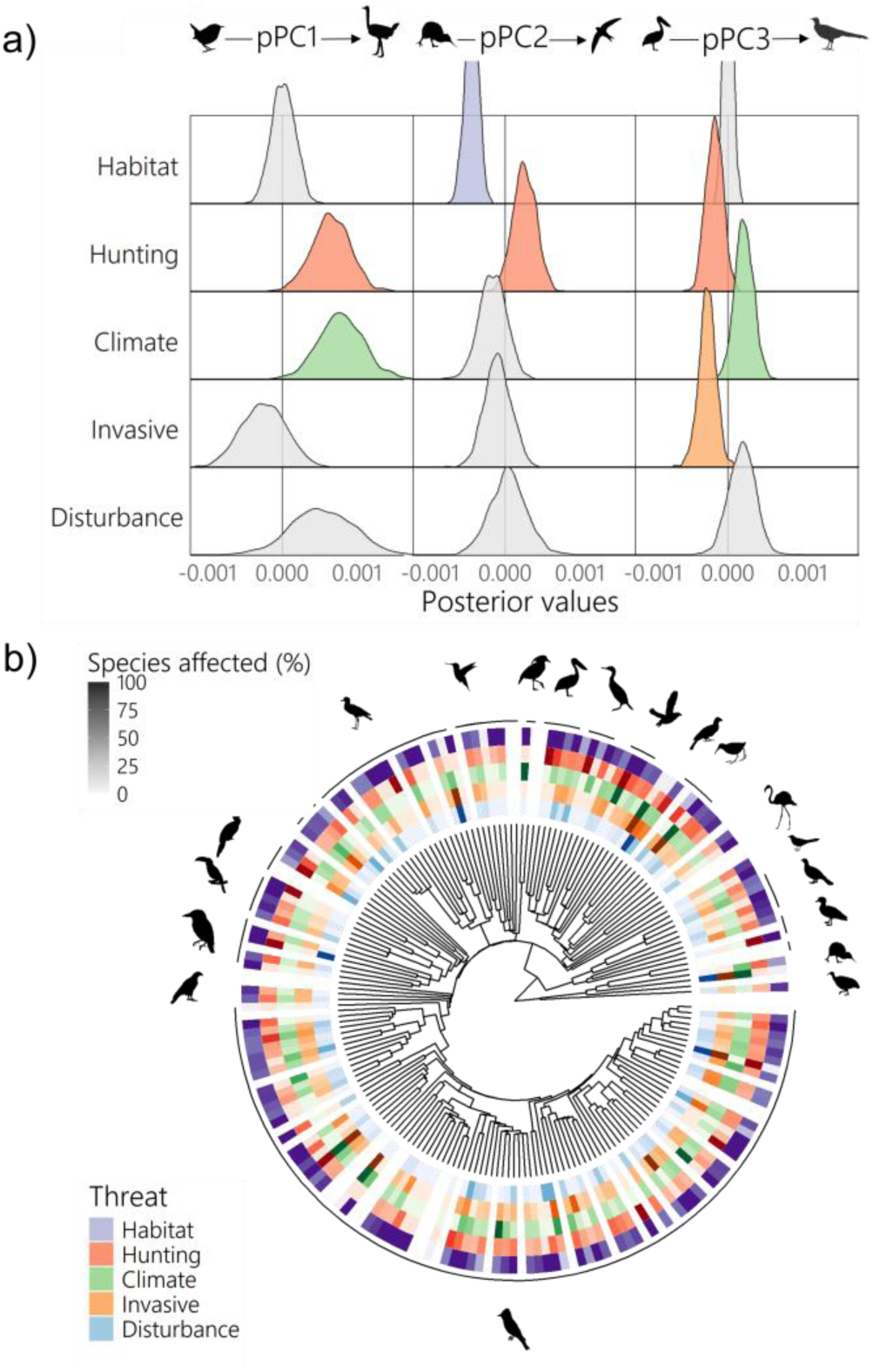
Drivers of extinction vary across morphospace and the avian tree of life. **a,** Posterior values from a multi-response Monte-Carlo generalized linear mixed model showing the relationships between each of the three phylogenetic principal component values and the frequency (from 1000 iterations) in which extinction was avoided under each driver-specific complete abatement scenario. pPC1 is a descriptor of body size, explaining variance in wing length, beak length (nares and culmen), beak depth, beak width, body mass, tail length, tarsus length and Kipp’s distance. pPC2 is associated with wing morphology, explaining variance in hand wing index and Kipp’s distance. pPC3 explains variance in beak length and shape and tail length. Least Concern species were not included in the extinction risk model as this is the lowest risk category so improvements under driver-specific complete abatement could not occur by definition. Least Concern species were assigned an extinction probability of 0.0001 over the next 100 years in all extinction scenarios (a low “background” extinction risk). **b,** Distribution of drivers of extinction with respect to phylogeny, shown by family (9873 species across 194 families, of which threat information is included for 2087 Near-Threatened and threatened species), with the intensity of colour reflecting the proportion of species in a family affected by each driver (threats affecting Least Concern species have not been comprehensively assessed so are not included, families including only Least Concern or Data Deficient species are shaded white). Hunting = hunting and collection; Climate= climate change and severe weather; Invasive = invasive species, genes and disease; Disturbance = disturbance and accidental mortality. Pollution was not included as it made a negligible contribution to functional richness loss (see Extended Data Table 2).

## Impact of six major drivers of extinction

Species’ traits shape their vulnerability to human activity, but different areas of trait space are affected by different threats^39,40^. As such, abatement of drivers of extinction could have differential outcomes for functional diversity. To test this, we focused on six drivers of extinction (Supplementary Dataset 1) and quantified the ‘*maximum avoidable contribution’* from each driver, describing the species and functional diversity loss avoided when the impact of current and future threats within each driver of extinction were completely removed, relative to diversity loss in the baseline scenario (see Methods). ‘Habitat loss and degradation’ had the highest maximum avoidable contribution, as driver-specific complete abatement was projected to avoid 1.4 ± 0.2% species richness loss (141 ± 24 species extinctions), and 0.9 ± 0.5% functional diversity loss (Fig. 1, Extended Data Table 2). Driver-specific complete abatement of ‘hunting and collection’ was projected to avoid 0.4 ± 0.5% functional diversity loss (Fig. 1), almost half that of ‘habitat loss and degradation’ despite requiring action for 591 species rather than 1658 species. Other drivers of extinction had smaller maximum avoidable contributions (Fig. 1). The relative magnitude of maximum avoidable contributions among drivers was comparable when simulating driver-specific partial abatement and driver-specific minimal abatement rather than driver-specific complete abatement.

While assessments of individual drivers on avian functional diversity exist^13,14,41^, assessments of multiple drivers simulatenously are rare, and are important for quantifying the relative impact of different drivers of extinction on avian functional diversity. We found that driver-specific complete abatement of ‘hunting and collection’ and ‘disturbance and accidental mortality’ was projected to result in disproportionately high avoidance of functional diversity loss for the number of species extinctions avoided (Extended Data Fig. 3). As abatement of different drivers of extinction had different value for the preservation of functional diversity, it is necessary to consider functional diversity in conservation planning and prioritisation.

To assess which species traits were vulnerable to drivers of extinction we used a mixed effects multi-response regression model of reduction in extinction risk under driver-specific complete abatement scenarios against values of three phylogenetic principal components (see Methods). Species could go extinct in 0 to 1000 of the iterations run for the baseline scenario, and species with a high certainty of high extinction risk had simulated extinction in more iterations than those with low extinction risk certainty or low extinction risk. If driver-specific complete abatement was effective at reducing extinction risk for a species, the number of iterations in which a species had simulated extinction was reduced. Reduction in extinction risk was therefore quantified as the number of iterations in which species extinction was avoided in driver-specific complete abatement scenarios, relative to the baseline scenario. A significant positive relationship indicates that abatement of a given driver of extinction reduces extinction risk in species with high values of a given phylogenetic principal component, and a significant negative relationship indicates that abatement of a given driver of extinction reduces extinction risk in species with low values of a given phylogenetic principal component (Fig. 3).

We project that the abatement of different drivers of extinction will benefit distinct morphologies. Birds with large body size (pPC1) were more likely to experience a reduction in extinction risk when abating ‘hunting and collection’ or ‘climate change and severe weather’ (Fig. 3, pMCMC<0.01 for both). While extinction risk bias towards species with large body size is widely reported^42,43^ we found that this was not the case for all threats, as there was no evidence of bias with respect to pPC1 for other drivers of extinction^39^.

The bias in extinction avoidance with respect to wing morphology (pPC2) was variable across drivers of extinction. Birds with broader wings (those with low pPC2 values) were more likely to experience a reduction in extinction risk under abatement of ‘habitat loss and degradation’ (pMCMC<0.01). In contrast, bird with slender wings (high pPC2 values) were more likely to experience a reduction in extinction risk under abatement of ‘hunting and collection’ (pMCMC=0.08, values under 0.1 treated as significant to give a one-tailed significance test of overlap with zero rather than default two-tailed test). Our finding that extinction avoidance was more likely for species with broad wings (low values of pPC2) when abating habitat loss and degradation was consistent with recent studies that show that birds with low hand-wing index (described by pPC2) are more sensitive to fragmentation^44^ and deforestation^45^.

The bias in extinction avoidance with respect to tail and beak morphology (pPC3) was also variable across drivers of extinction. Species with long tails and short beaks (low pPC3 values) were more likely to experience a reduction in extinction risk following abatement of both ‘hunting and collection’ and ‘invasive species, and diseasè (pMCMC=0.96 and pMCMC<0.05 respectively [hunting and collection not significant when insignificant variables removed]), whilst species with short tails and long beaks (high pPC3 values) were more likely to experience a reduction in extinction risk following abatement of ‘climate change and severe weather’ (pMCMC=0.08). Reduced extinction risk following abatement of climate change and severe weather was associated with traits involved in thermoregulation^46–49^. Birds with large body size (pPC1), but also large beaks (pPC3) were more likely to avoid extinction when climate change and severe weather was abated. As a bird’s beak also influences their trophic niche^50^, failing to mitigate species decline due to climate change could have knock-on effects for trophic interactions. Variable extinction avoidance across functional trait space suggests that prioritising threat abatement based on the magnitude of projected biodiversity loss alone is inappropriate. Reducing the impact of multiple drivers of extinction is necessary to ensure that diverse functional morphologies are conserved.

## The potential of targeted species recovery programmes

Even with ambitious action, large-scale threat abatement will not prevent all species extinctions and functional richness loss in the next 100 years. As such, targeted species recovery programmes will be needed, which we define as in-situ and ex-situ measures to boost species survival and reproductive success that do not involve threat reduction. Here we explore one possible approach, quantifying the benefits of preventing a small number of species extinctions targeted to reduce the loss of global functional richness. Using a metric of functional uniqueness which describes the probability of functional richness loss due to species extinction, we identified the most unique threatened species among the 9873 bird species studied. Preventing the extinction of the most unique species (we tested scenarios protecting between 40 and 200 species, Extended Data Fig. 4), was effective at reducing projected functional diversity loss. We found that preventing extinction of the top 100 most unique threatened species avoided 68 ± 5% of projected functional diversity loss under the baseline scenario compared to the 26 ± 13% avoided by partial abatement of all threats for all species. By preventing the extinction of 100 species (1% of species), 2.2 ± 0.32% of functional diversity could be conserved, if the most functionally unique threatened species were prioritised. This approach would require the avoidance of 37 ± 25 projected extinctions in the next 100 years. Bolam et al.^17^ found that 21-32 bird species have been saved from extinction by conservation efforts since 1993, suggesting that this could be an achievable goal (although 10 extinctions occurred despite management).

The most functionally unique birds span taxonomic and ecological groups, from the Sulu Hornbill (*Anthracoceros montani*) of the southernmost Philippine islands^51^, to the Ascension Frigatebird (*Fregata aquila*) that patrols the Atlantic Ocean. Some are wide-ranging like the Southern Royal Albatross (*Diomedea epomophora*), whilst others are thought to only survive in one location, like Stresemann’s Bristlefront (*Merulaxis stresemanni*). Unique species include scavengers, such as the Andean condor (*Vultur gryphus*), nectarivores, such as the Yellow-bellied sunbird-asity (*Neodrepanis hypoxantha*), vertivores, such as the Madagascar Serpent-eagle (*Eutriorchis astur*), and frugivores such as the Bare-necked umbrellabird (*Cephalopterus glabricollis*). A full list of the top 200 most unique threatened birds is given in Supplementary Dataset 2.

Previous studies have found that functionally unique species are more likely to be threatened with extinction than less functionally unique species^30,52^. We provide evidence that conservation strategies for birds should prioritise functionally unique species, as has been proposed for other taxonomic groups^53,54^. Besides their inherent value, functional unique species are more likely to be used by humans, for food, material and medicine, so preventing the extinction of functionally unique species could be important for the delivery of ecosystem services^55^. Effective targeted recovery programmes that explicitly consider species uniqueness hold great potential for conserving global functional diversity as a complementary strategy to threat abatement.

## Conclusions

Both large-scale protection from the drivers of extinction and targeted species recovery programmes will be needed to prevent avian extinctions and functional diversity loss in the next 100 years. While not effective at preventing all biodiversity loss, threat-abatement is essential for ensuring that species that currently have healthy, stable populations, do not go into decline^15^. Nevertheless, our findings suggest that conservation policy should not focus solely on large-scale protection from the drivers of extinction, as even in ambitious scenarios, only half of projected species extinctions and functional diversity loss attributable to these drivers of extinction could be avoided.

Reducing the impact of different drivers of extinction protected distinct areas of functional trait space. Abatement of ‘habitat loss and degradation’ made the greatest overall contribution to avoided species extinctions and functional diversity loss, but management of ‘hunting and collection’ and ‘disturbance and accidental mortality’ prevented greater functional diversity loss proportional to the number of species projected to go extinct. Because different areas of functional trait space were impacted by different drivers, abatement of all drivers of extinction is necessary to conserve functional diversity.

When completely or partially abating the drivers of extinction, functional richness loss was correlated with species extinctions, so reducing species extinctions is projected to reduce functional diversity loss. However, targeted species recovery programmes that focus on functionally unique species hold great potential for the conservation of functional diversity, whilst requiring conservation of relatively few species. By conserving the top 100 most unique threatened species it may be possible to prevent more than 2/3 of the projected functional diversity loss through avoiding ∼37 species extinctions. While prioritisation of recovery programmes offers great potential for protecting functional diversity, the ethical questions about prioritising some species over others, and the risks of overlooking ecosystem functions and services provided by other species, whether known or unknown, must be considered. If human activity continues to affect biodiversity as it does today, we project that in the next 100 years we will lose more than three times the number of bird species as has been lost since 1500. It is therefore urgent that we decide which dimensions of biodiversity we wish to protect, consolidate their measurement, and include them into every stage of conservation planning, monitoring, and impact assessment.

## Methods

### Data collection

We used data on species morphological and geographical traits, threats and phylogenetic relationships, to conduct this study. Trait data for 11003 extant bird species^56^ were obtained from AVONET^26^. Under the BirdLife taxonomy, only 8% of species in AVONET have imputed data for one or more traits, and <5% of species have imputed data for more than one trait. For all study species, data on threats were obtained in June, 2022 from the IUCN Red List^20^ using the function *rl_threats* in the package *rredlist*^57^ in R^58^. Bird species are reassessed every four years so there can be a slight delay between species decline or recovery and reported change in extinction risk category^59^. Taxonomic discrepancies between AVONET and IUCN threat data for 141 species were reconciled using the function *rl_synonym*. One-to-one matches were found for all species so these taxonomic differences will not impact the results.

Phylogenetic data were obtained from Jetz et al.^60^. A maximum clade credibility tree was constructed from the first 1000 trees based on the Hackett backbone^60,61^. Jetz et al.^60^ included 9993 species in their analysis; we refer to the species nomenclature and taxonomic treatments adopted in this study as “BirdTree taxonomy”. A matching procedure to translate between BirdLife and BirdTree taxonomies was needed to enable analysis of functional diversity loss whilst accounting for phylogenetic covariance between species. Differences between the BirdLife^56^ and BirdTree^60^ taxonomy were reconciled using the crosswalk provided with AVONET^26^ (see Supplementary information). This gave 9879 selected synonym matches between BirdLife and BirdTree (89.9% of BirdLife synonyms and 98.9% of BirdTree synonyms). Repeating the analysis with all BirdLife synonyms and with principal component analysis that did not account for phylogenetic covariance between species, had a small impact on the percentage of projected species extinctions and functional diversity loss but did not affect our conclusions (see Supplementary information). Five species treated as Extinct in the Wild and one species listed as Extinct by IUCN^20^ but not listed as Extinct in the AVONET crosswalk (*Zosterops conspicillatus*) were removed from the analysis giving a total of 9873 species.

### Estimating functional diversity

Functional diversity quantifies the diversity of functional traits within an assemblage, defined as the measurable characteristics of an organism which influence its ecological niche^62,63^. We used 11 continuous morphological traits extracted from AVONET^26^, including body mass, and linear measurements of beak, wing, tail and tarsus. These traits collectively capture bird ecological niches through their association with diet, dispersal and habitat^25–27^. Using continuous morphological traits enables more fine-grained discrimination between bird species sharing the same ecological groups, thus providing more in-depth information about ecological variation between species than categorical ecological traits^50^. As life-history traits are more useful for explaining variation in species response to human activity rather than the ecological impacts of decline^26^, they were not included in functional diversity estimations. Trait data were log10 transformed and scaled to unit variance.

We used phylogenetic principal component analysis (pPCA) to reduce dimensionality. Principal component analysis produces axes that are mathematically uncorrelated, but axes may be phylogenetically correlated if species trait data are non-independent due to shared evolutionary history^64,65^. pPCA accounts for phylogenetic correlation between axes by removing phylogenetic covariance and calculating major axes of non-phylogenetic residual variation^65^. pPCA was carried out using the *phyl.pca* function in *phytools*^66^, based on covariance and using lambda to obtain the correlation structure which was optimised using Restricted Maximum Likelihood (REML). The first three phylogenetic principal components (pPCs) described over 80% of variance in the dataset (87.2%), and adding more pPCs described comparatively less variation (identified through scree plots, Extended Data Fig. 5). We therefore used the first three pPCs to summarise variation in the dataset (Extended Data Tables 4 and 5). Using the first two or four pPCs instead of three and using alternative ordination methods did not affect our conclusions (see Supplementary Information).

We calculated global functional richness for the whole assemblage (9873 species) using trait probability densities^28^. Firstly, a multivariate gaussian probability distribution was fitted for each species (Extended Data Fig. 6), where the mean was provided by pPC values derived from functional trait data^26^ and standard deviation was estimated using a bandwidth selector (*Hpi.diag* function from package *ks*^67^). Next, we took the sum of species probability distributions to give the community trait probability density (Extended Data Fig. 6). This was implemented through the *TPDsMean* and *TPDc* functions in package *TPD*^68^, with 50 divisions for each pPC (meaning that trait space was divided into 125 000 grid cells). We calculated functional richness using the *REND* function in package *TPD*, which calculates the volume of trait space occupied by the community trait probability densities.

### Modelling extinction risk

To predict how threat reduction affects projected avian diversity loss we constructed a model of species extinction risk (IUCN Red List category^20^) with threats as explanatory variables and accounting for spatial and phylogenetic covariance (referred to as ‘the extinction risk model’). The extinction risk model allowed us to quantify the independent contribution of each threat to extinction risk despite the fact that many species were affected by multiple threats (2542 species out of 9873, not including Extinct in the Wild and Extinct species), while comparatively few were affected by only one (638 out of 9873 species). Overlooking non-independence between threats can result in misleading findings about the relationship between threats and extinction risk and patterns of bias in impacts of these drivers across species assemblages^69^. Threats varied across the tree of life (Fig. 3) and spatially (Supplementary Fig. 1) so we included phylogeny and spatial variables as random effects.

The extinction risk model was fitted using a Markov chain Monte Carlo multivariate generalised linear mixed model (MCMCglmm) to predict extinction risk for species that were listed by the IUCN Red List as Near Threatened, Vulnerable, Endangered, and Critically Endangered. Threats have been comprehensively described for 99% of species in these categories. MCMCglmms were fitted using the R package *MCMCglmm*^70^. We used 39 pseudo-continuous fixed effects, describing the expected percentage population decline over a 10-year period or three generations (from threat scope and severity data, see details below and Extended Data Table 3) due to each threat under the second level classification described by the IUCN^71^ (for example, “*1.1 Housing & Urban Areas*” and “*1.2 Commercial and Industrial Areas*”). Threats which affected 10 or fewer species were grouped with other threats. “*4.3 Shipping lanes*” and *“4.4 flight paths*” were grouped with “*4.1 roads and railroads*”. “*10.1 Volcanoes*”, “*10.2 Earthquakes/tsunamis*” and “*10.3 avalanches/landslides*” were grouped together. “*8.3 Introduced genetic material*” and “*8.6 Disease of unknown cause*” were grouped with “*8.4 problematic species/diseases of unknown origin*”.

Threats were assigned an expected population decline (over a 10-year period or three generations) based on their scope (percentage of species range affected by a threat) and severity, following Garnett et al.^72^ and Mair et al.^33^ (Extended Data Table 3). Where there were multiple threats listed for the same threat category under the second order classification listed by the IUCN^71^ the maximum expected population decline was used. For example, *Acrocephalus familiaris* is experiencing slow, continuous declines due to the invasive species *Schistocerca nitens* across the majority of its range, but it is also experiencing rapid declines due to *Oryctolagus cuniculus* across the whole of its range. For “*8.1 Invasive non-native/alien species/diseases*”, *Acrocephalus familiaris* was assigned an expected population decline of 24% associated with rapid declines across the whole of its range (Extended Data Table 3). We focused on second order threats rather than third order threats because threats are not always specified at the third order level. Threats which were expected to cause no decline or negligible declines across a majority or minority of a species range, had an expected population decline of 0, so were effectively discarded.

Across all species, 11.58% of threat data were missing scope or severity values. Missing scope and severity data were imputed with missForest imputation (implemented through R package *missForest*^73^) from threat type, scope, severity and timing, and incorporating phylogenetic data through eigenvectors^74^. Removing a similar proportion of values from complete data on threat timing, scope and severity to test imputation accuracy gave a mean accuracy of 82.5% (see Supplementary information). 48 rows were also missing data on timing (needed for creating extinction scenarios, see “*Extinction scenarios”* section of the method), and these data were imputed in the same way as scope and severity.

Initially, phylogeny, minimum latitude, maximum latitude, centroid latitude and centroid longitude were included in the model as random effects (see Supplementary information for more details), where centroid latitude and centroid longitude described the midpoint of species ranges. Spatial variables were obtained from AVONET and had been calculated from species’ breeding and resident ranges including areas where the species was coded as extant and either native or reintroduced^26^. We determined if random effects were useful for explaining variation in extinction risk by plotting posterior random effect variances (Supplementary information Fig. 1). The posterior random effect variances of centroid latitude clustered close to zero, suggesting that the random effect was not informative^75^. As such, centroid latitude was not included as a random effect in the final extinction risk model. Seventeen species were not included in the extinction risk model as they had incomplete spatial information. Species with missing spatial information were either Possibly Extinct, had no known breeding or resident range or their range data had been redacted to protect them from trafficking risk^26^. Twenty-two species had no threat data listed and were not included in the extinction risk model. The final model was fitted for the remaining threatened and Near-Threatened species (*n*=2087).

For fixed effects, Cauchy-scaled Gelman priors were used (with an expected value of 0), as is recommended by Gelman^76^ for ordinal regressions. For the phylogenetic random effect, we used a chi-squared prior (expected covariance of 1, degree of belief of 1000, mean vector of 1 and covariance matrix of 1), as this best approximates a uniform distribution, giving an uninformative prior^70,77,78^. For spatial random effects we used parameter expanded priors (expected covariance of 1, degree of belief of 1, mean vector of 0 and covariance matrix of 625) as they are often less informative than the default inverse-Wishart prior^79^. Because it is not possible to estimate the residual variance with an ordinal response variable (extinction risk), the residual variance was fixed to 1 following Hadfield (2017)^79^. The model was insensitive to changes in prior specification (see Supplementary information). MCMC chains were run for 103000 iterations, with a burn in period of 3000 iterations and a thinning interval of 100 iterations. Model convergence was assessed using plots of parameter traces produced through the *plot* function in package *MCMCglmm*^70^. Non-significant fixed effects were removed iteratively, removing the least significant fixed effect, rerunning the model and removing the least significant fixed effect again, until only significant fixed effects remained. Significance was assessed using the pseudo p value (*“pMCMC*”), estimated by *MCMCglmm*^70^. The pseudo p value is calculated by finding the probability that the posterior is greater or less than zero, whichever is smaller, and multiplying by two^80^. A significance threshold of 0.1 rather than 0.05 was used, to give a one-tailed significance test that the posterior distribution overlaps with zero, rather than the default two-tailed significance test. The final model structure was,

Extinction risk category ∼ X1.2 + X1.3 + X2.1 + X2.2 + X2.3 + X4.2 + X5.1 + X5.3 + X5.4 + X6.3 + X7.1 + X7.2 + X8.1 + X8.1 + X8.2 + X9.3 + X10.1 + X11.1 + X11.4 + X12.1 + random (phylogeny + minimum latitude + maximum latitude + centroid longitude),

Where X1.2 refers to the expected percentage population decline over a 10-year period or three generations (Extended Data Table 3) as a result of IUCN second order threat “*1.2 Commercial and Industrial Areas”* and so on. X10.1 includes threat impact from X10.1, X10.2 and X10.3 as the threats were grouped together because each threat affected less than ten species (see Supplementary Dataset 1 for threat codes, threat descriptions and model parameter estimates).

Model accuracy was assessed by calculating the proportion of species for which the category listed by the IUCN^20^ matched the category that most frequently (across iterations) had the highest probability. Model accuracy was tested through self-prediction, rather than out of bag prediction as random effects were important for explaining variance in extinction risk, and we did not intend on using the model on species which were not in the model data.

### Projected diversity loss

We used the extinction risk model to predict the probability that species belonged to each Red List category, then using expected probability of extinction for each Red List category, simulated extinctions likely to occur in the next 100 years. We used an approach that explicitly incorporated uncertainty in model estimates and stochasticity in realised extinctions given extinction probabilities.

The extinction risk model returned 1000 posterior estimates per species (1000 iterations) of the probability that the species belonged to each extinction risk category (Near Threatened, Vulnerable, Endangered and Critically Endangered). Posterior estimates were extracted from the model using the function *predict2* from the *postMCMCglmm* package^81^. All Least Concern species and 39 Near Threatened or threatened species that were missing spatial or threat data (species not included in the extinction risk model), were assigned a probability of 1 of belonging to their Red List category as currently listed by the IUCN. Species classified as Data Deficient (n=41) were conservatively assigned to the Least Concern category.

An overall probability of extinction was then calculated for each species according to equation 1.

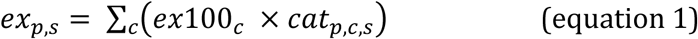

Where *ex*_*p*,*s*_is the probability that species *s* goes extinct in the next 100 years according to the posterior estimation *p*; *ex*100_*c*_is the assigned probability that a species in IUCN extinction risk category *c* goes Extinct in the next 100 years, and *cat*_*p*,*c*,*s*_ is the probability that species *s* belongs to IUCN extinction risk category *c* according to posterior estimation *p* or, for the species not included in the extinction risk model, a probability of 1 for their Red List category as currently listed by the IUCN, and a probability of 0 for all other Red List categories. *ex*100_*c*_were based on previous work^24,29,82^ and set to: Critically Endangered = 0.999, Endangered = 0.667, Vulnerable = 0.1, Near Threatened = 0.01 and Least Concern = 0.0001.

Estimates of *ex*_*p*,*s*_were converted to a binary outcome of extinct or extant using the R function ‘*sample*’, where the probability of being assigned extinct was *ex*_*p*,*s*_. For each scenario we report the mean and standard deviation in the number of extinctions across 1000 iterations as a percentage of the number of species for which extinctions were simulated (9873 species). Functional diversity loss was estimated by removing species projected to go extinct, calculating functional diversity across species predicted to be extant and comparing to functional diversity of the full assemblage (9873 species).

### Threat reduction scenarios

We estimated projected loss in species and functional diversity in the next 100 years under a baseline extinction scenario, and three threat reduction scenarios: complete abatement, partial abatement and minimal abatement. Under the baseline scenario, the extinction risk model was used to predict the probability that species belonged to Red List categories, assuming that the impact of all threats remains as is currently listed by IUCN^20^, following the method for predicting extinctions as detailed in the section above. Under the complete abatement scenario predictions from the extinction risk model were obtained after setting the expected population decline to zero for all threats with a timing of “*Ongoing”*, “*Past, Likely to Return”* and “*Future”*. The expected population decline of threats with a timing of *“Past, Unlikely to Return”* was not changed as they cannot be prevented but could still contribute to the extinction risk of a species through extinction lags (although predicted extinctions were similar if expected population decline for threats with a timing of “*Past, Unlikely to Return*” was set to zero, see Supplementary information). Under the partial abatement scenario threat impacts were altered to simulate removal of threats across at least 50% of species ranges by reassigning expected population declines for threats with a scope of “*Whole (>90%)*” or “*Majority (50-90%)*” to the decline expected for a scope of “*Minority (<50%)*” (Extended Data Table 3). Under the minimal abatement scenario threat impacts were altered to simulate removal of threats across at least 10% of species ranges by reassigning expected population declines of threats with a scope of “*Whole (>90%)*” to expected decline for a scope of “*Majority (50-90%)*”. Least Concern species were not included in the extinction risk model as this is the lowest risk category so could not show a reduction in extinction risk under abatement scenarios by definition.

We used Cohen’s D to quantify the effect size of the difference in means of diversity loss under each threat reduction scenario divided by their pooled standard error. p values were not reported as the sample size (number of iterations) could be increased easily, reducing standard error and giving significance even when there was a very small differences in means, leading to type 1 errors.

### Vulnerable bird morphologies and hotspots of conservation potential

To find which bird morphologies had greatest extinction risk, we plotted loss in density of trait space occupation under the baseline scenario using trait probability densities. We constructed a community probability distribution by taking the sum of species probability distributions, where each species probability distribution was given a weight between 0 and 1000 describing the number of iterations in which they did not go extinct under the baseline scenario. The resulting community probability distribution was rescaled to show absolute rather than relative change in density of occupation of trait space. For each cell we compared the density of occupation of the whole assemblage to the density of occupation in iterations of the baseline extinction scenario. The loss in density was calculated as a percentage of the density of trait space occupation in the full assemblage where all species had a value of 1000 indicating no extinction (Extended Data Fig. 7). Areas of trait space with high loss in density (close to 100%) are predicted to have a high risk of extinction.

To find which bird morphologies had greatest reduction in extinction risk under complete abatement of all threats (protection in Extended Data Fig. 7), we plotted the density of trait space occupation under the complete abatement scenario using trait probability densities, where all species were given a weight according to the number of iterations in which extinction was avoided in the complete abatement scenario relative to the baseline scenario. For plotting loss in density under the baseline scenario and loss in density averted under the complete abatement scenario, phylogenetic principal components were divided into 100 bins to give high plot resolution. To facilitate visualisation, this was carried out in two dimensions (two phylogenetic principal components at a time).

### Impact of six major drivers of extinction

We aimed to assess the independent contribution of six major drivers of extinction to projected avian diversity loss. Threats in the extinction risk model were grouped into six “*drivers of extinction*”: ‘habitat loss and degradation’, ‘hunting and collection’, ‘climate change and severe weather’, ‘disturbance and accidental mortality’, ‘invasive species and diseasè and ‘pollution’ (for threat classification see Supplementary Dataset 1). Geological events and threats described as “*Other*” were grouped into an ‘other’ category. Whilst these ‘other’ threats were included in the model of extinction risk and their impact was accounted for when assessing the impact of all drivers of extinction together, we did not assess their impact individually.

We projected species and functional diversity loss under driver-specific complete abatement scenarios where the impact of threats in a given driver of extinction with a timing of “*Ongoing*”, “*Past, Likely to Return*” and “*Future*” was removed by setting their expected population decline to zero. The “*maximum avoidable contribution*” was calculated as the difference in predicted species and functional diversity loss between the baseline scenario and the driver-specific complete abatement scenario.

We used Cohen’s D to quantify the effect size of the difference in means of diversity loss under the baseline and driver-specific complete abatement scenarios divided by their pooled standard error. As before, p values were not reported as the sample size (number of iterations) could be increased easily, leading to type 1 errors.

To find the severity of functional richness loss under a given driver of extinction in relation to the number of species projected to go extinct, we used a linear mixed effects model to describe the functional richness loss avoided, using the number of species extinctions avoided, and the driver of extinction (categorical) as explanatory variables. Model iteration was used as a random effect to account for non-independence in the calculated differences in functional diversity loss and species extinctions from baseline to threat reduction scenarios. The dredge function from the *MuMin* package^83^ was used to identify the best model from all combinations of explanatory variables and the two-way interaction between species extinctions and driver of extinction. The best model included the number of species extinctions avoided and the driver of extinction, but not the interaction between species extinctions avoided and the driver of extinction.

### Biases in extinction avoidance

We aimed to find whether abatement of different drivers of extinction could avoid extinction in different regions of trait space. We constructed a multi-response Markov-Chain Monte-Carlo mixed effects model, with the three phylogenetic principal component values for each species as response variables (including all 9873 species studied), and the number of iterations in which extinction was avoided under driver-specific complete abatement for each driver of extinction as explanatory variables, accounting for phylogenetic covariance. Pollution was not included as an explanatory variable as driver-specific complete abatement of pollution had a negligible impact on functional richness loss. The residual structure was allowed to vary for each response variable. Random effect priors were provided as a diagonal matrix, with an expected covariance between response variables of 0, and expected variance within response variables of 1. The degree of belief parameter for the random effect prior was 2 ^84^. The expected value of fixed effects and the theta-scale parameter were 0, with a covariance matrix where the expected covariance between fixed effects was 0 and the expected variance within fixed effects was 1 x 10^10^. Posterior distributions were plotted and a significance threshold of 5% overlap with 0 was used (pMCMC=0.1).

### The potential of targeted species recovery programmes

To identify the potential value of preventing the extinction of the most functionally unique threatened species we calculated functional uniqueness for all species (Extended Data Fig. 8). For each species we calculated the proportion of the density of the community probability distribution that was occupied by the species probability distribution, for each grid cell where the species probability density was greater than 0. We calculated the mean proportion for each species across grid cells where the species probability distribution was greater than 0, giving greater weight to cells where the species distribution had greater probability (higher density). The maximum uniqueness possible was 1, indicating that a species probability distribution had no overlap with the probability distribution of any other species included in the analysis. Uniqueness tended towards a lower limit of 0, indicating that the species probability distribution had high overlap with many other species probability distributions probability distributions, and many other species occupied the same area of trait space.

We then identified the most unique and threatened (listed as Vulnerable, Endangered or Critically Endangered) species as potential targets for action. We compared functional richness loss in the baseline scenario, with functional richness loss when extinctions of the most unique threatened species were prevented. We explored the consequences of avoiding extinction for the top 40 to 200 most unique threatened species (in intervals of 20 species). 1000 posterior estimates were obtained for each extinction scenario.

## Acknowledgements

Kerry Stewart acknowledges PhD studentship funding from the SCENARIO NERC Doctoral Training Partnership grant NE/S007261/1. Chris Clements was funded by the Natural Environment Research Council (grants NE/Z000130/1 and NE/T006579/1). Carlos P. Carmona was funded by the Estonian Research Council (grants PRG2142 and MOBERC100) and the European Union (ERC, PLECTRUM, 101126117).

## Data and code availability

AVONET data on morphological, ecological and geographical traits for all birds^26^ is available for use under the creative commons licence (CC BY 4.0): https://doi.org/10.6084/m9.figshare.16586228.v7. Data on IUCN extinction risk categories, and threats affecting each species, are available from the IUCN Red List^20^ and can be accessed through the package *rredlist*^57^. Information on terms of use of IUCN Red List data can be found at https://www.iucnredlist.org/terms/terms-of-use. Code used for figures and analyses and supplementary datasets are presented in https://figshare.com/s/855f64d2608140f295ab (url for a public repository will be made available on acceptance).

## Extended Data

**Extended Data Table 1.**
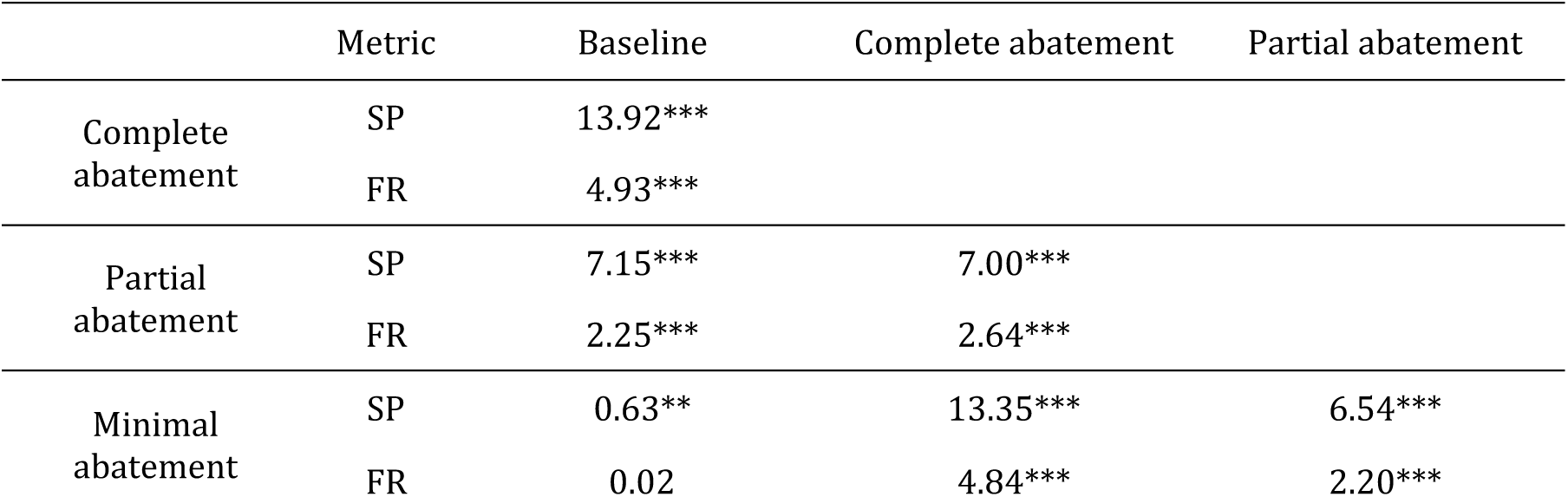
Predicted effect (Cohen’s D) of management scenarios on diversity loss (1000 iterations for each extinction scenario. ,SP = number of species extinctions and FR= percentage functional richness loss, * = small to medium impact [0.2 < Cohens D < 0.5], ** = medium to large impact [0.5 < Cohens D < 0.8], *** = large impact [Cohens D > 0.8]^85^).

**Extended Data Table 2.** Predicted effect (Cohen’s D) of driver-specific complete abatement (1000 iterations for each extinction scenario. ,\* = small to medium impact [0.2 < Cohens D < 0.5], ** = medium to large impact [0.5 < Cohens D < 0.8], *** = large impact [Cohens D > 0.8]^85^).

| Driver of extinction | Species richness | Functional richness |
| --- | --- | --- |
| Habitat loss and degradation | 7.37*** | 2.29*** |
| Hunting and collection | 2.23*** | 1.04*** |
| Climate change and severe weather | 1.98*** | 0.35* |
| Invasive species and disease | 1.96*** | 0.42* |
| Disturbance and accidental mortality | 0.83*** | 0.49* |
| Pollution | 0.44* | 0.10 |

**Extended Data Table 3.** Expected population decline over a 10-year period or three generations (%) from scope and severity scores (from Garnett et al.^72^ and Mair et al.^33^). Scope describes the percentage of a species range covered by a threat.

|  |  | Severity |  |  |  |  |  |
| --- | --- | --- | --- | --- | --- | --- | --- |
|  |  | Very rapid declines | Rapid declines | Slow, significant declines | Negligible declines | No decline | Causing/could cause fluctuations |
| Scope | Whole (>90%) | 63 | 24 | 10 | 1 | 0 | 10 |
|  | Majority (50-90%) | 52 | 18 | 9 | 0 | 0 | 9 |
|  | Minority (<50%) | 24 | 7 | 5 | 0 | 0 | 5 |

**Extended Data Table 4.** Variance explained (%), cumulative variance explained (%), and eigenvalues of phylogenetic principal components (pPC) (*n*=9873 species).

|  | Variance explained (%) | Cumulative variance explained (%) | Eigenvalues |
| --- | --- | --- | --- |
| pPC1 | 62.8 | 62.8 | $1.78 \times 10^{-2}$ |
| pPC2 | 15.5 | 78.4 | $4.39 \times 10^{-3}$ |
| pPC3 | 8.9 | 87.2 | $2.50 \times 10^{-3}$ |
| pPC4 | 4.1 | 91.3 | $1.15 \times 10^{-3}$ |
| pPC5 | 3.9 | 95.2 | $1.10 \times 10^{-3}$ |
| pPC6 | 2.1 | 97.2 | $5.84 \times 10^{-4}$ |
| pPC7 | 1.1 | 98.4 | $3.15 \times 10^{-4}$ |
| pPC8 | 0.9 | 99.3 | $2.54 \times 10^{-4}$ |
| pPC9 | 0.5 | 99.7 | $1.27 \times 10^{-4}$ |
| pPC10 | 0.2 | 99.9 | $5.49 \times 10^{-5}$ |
| pPC11 | 0.1 | 100.0 | $2.92 \times 10^{-5}$ |

**Extended Data Table 5.** Eigenvectors of phylogenetic principal components (pPC) (*n*=9873 species).

|  | pPC1 | pPC2 | pPC3 | pPC4 | pPC5 | pPC6 | pPC7 | pPC8 | pPC9 | pPC10 | pPC11 |
| --- | --- | --- | --- | --- | --- | --- | --- | --- | --- | --- | --- |
| Body mass | 0.29 | -0.04 | -0.02 | -0.20 | -0.23 | -0.11 | 0.72 | 0.34 | 0.37 | 0.16 | -0.08 |
| Wing length | 0.32 | 0.17 | 0.20 | -0.09 | -0.18 | 0.45 | -0.04 | 0.00 | -0.01 | -0.61 | -0.47 |
| Tarsus length | 0.28 | -0.15 | 0.07 | -0.13 | -0.63 | -0.55 | -0.41 | -0.12 | 0.05 | -0.02 | -0.02 |
| Beak length (culmen) | 0.35 | -0.09 | -0.41 | 0.48 | 0.09 | 0.17 | -0.19 | -0.14 | 0.60 | -0.05 | 0.13 |
| Beak length (nares) | 0.36 | -0.11 | -0.37 | 0.44 | -0.07 | -0.07 | 0.16 | 0.16 | -0.67 | 0.06 | -0.16 |
| Beak width | 0.32 | -0.05 | -0.23 | -0.48 | 0.44 | -0.08 | -0.38 | 0.52 | -0.01 | 0.00 | -0.01 |
| Beak depth | 0.31 | -0.09 | -0.18 | -0.39 | 0.29 | -0.11 | 0.24 | -0.74 | -0.11 | -0.02 | -0.03 |
| Hand-wing index | 0.02 | 0.78 | -0.15 | 0.03 | 0.01 | -0.15 | -0.11 | -0.08 | 0.09 | 0.38 | -0.42 |
| Secondary1 | 0.34 | -0.03 | 0.26 | -0.13 | -0.20 | 0.56 | -0.16 | -0.07 | -0.12 | 0.60 | 0.22 |
| Kipps Distance | 0.19 | 0.55 | -0.05 | -0.04 | -0.11 | -0.04 | 0.10 | 0.04 | -0.14 | -0.32 | 0.72 |
| Tail length | 0.37 | 0.03 | 0.69 | 0.34 | 0.42 | -0.32 | 0.03 | 0.01 | 0.03 | 0.02 | 0.00 |

**Extended Data Figure 1.**
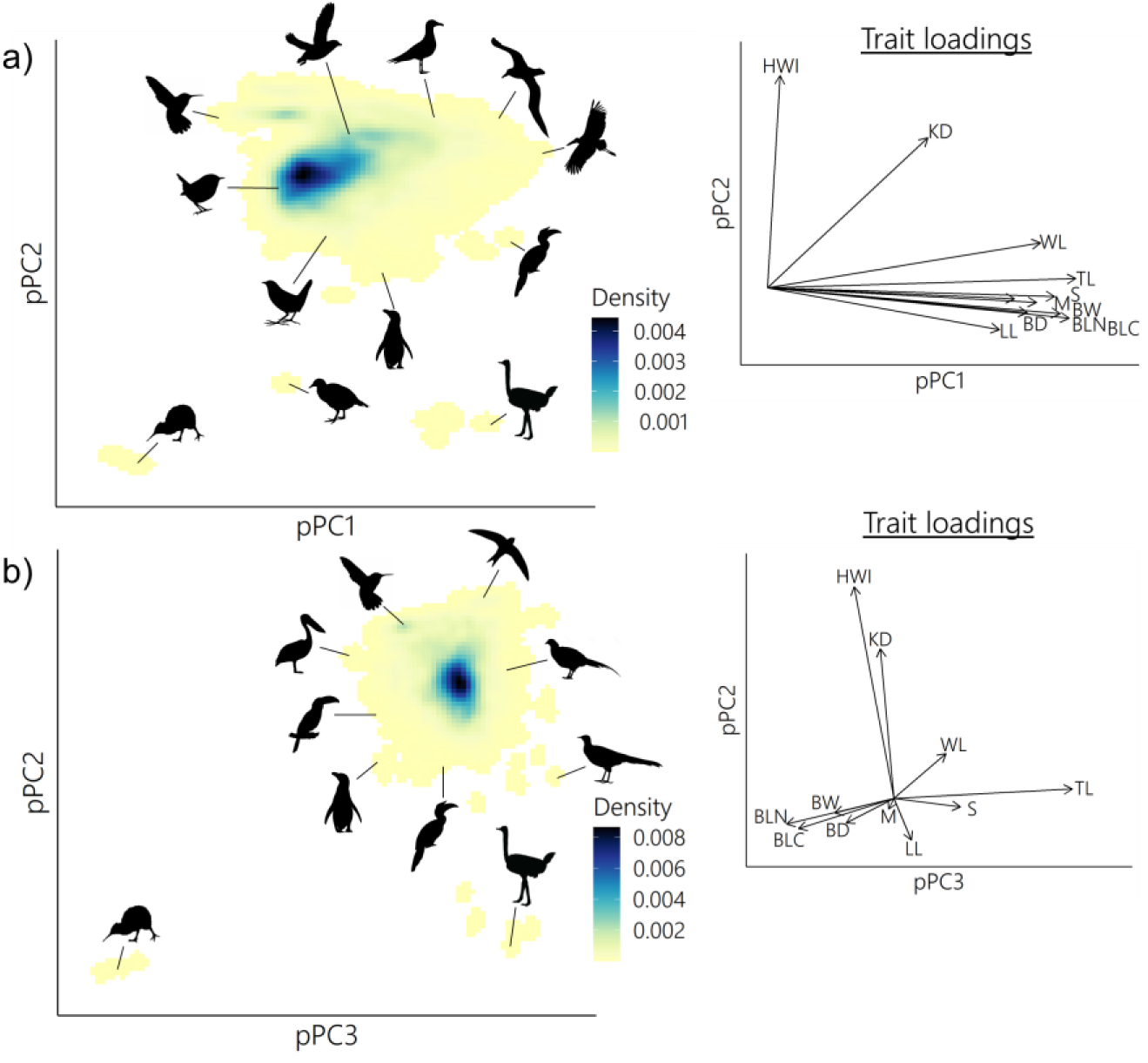
Occupation of morphospace in extant birds. Shown on 2-dimensional plane with respect to **a,** pPC1 and pPC2, and **b,** pPC2 and pPC3 with trait loadings (also see Extended Data Table 5). BD = beak depth, BLC = beak length (culmen), BLN = beak length (nares), BW =beak width, HWI = hand-wing index, KD = Kipp’s distance, LL = tarsus length, M = body mass, S = first secondary length, TL = tail length, WL = wing length (*n*=9873 species).

**Extended Data Figure 2.**
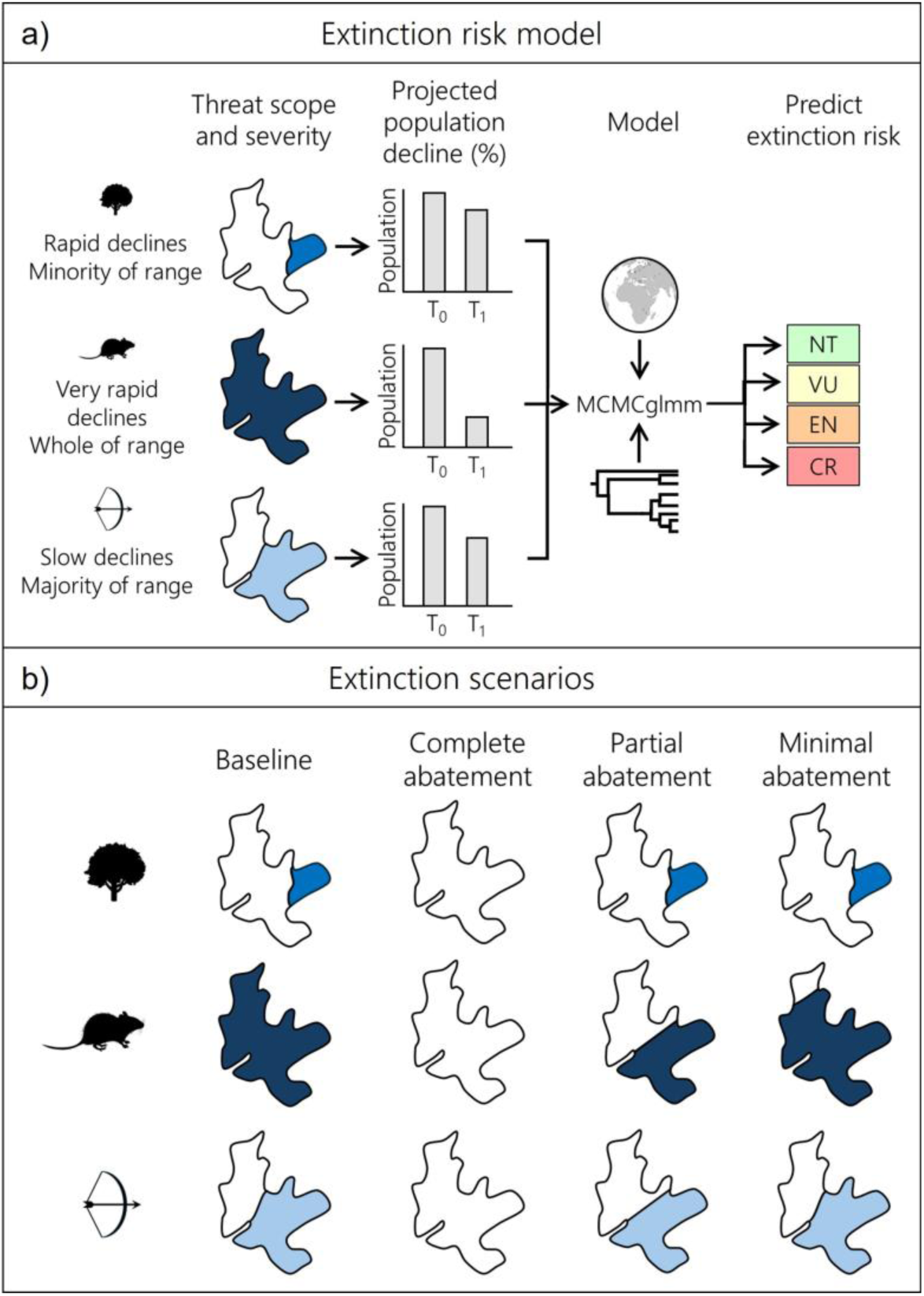
The extinction risk model and extinction scenarios. **a,** IUCN Red List data^20^ on threat scope and severity were used to assign projected population decline over a 10-year period or three generations according to Garnett et al.^72^ and Mair et al.^33^. Data on projected population decline for all species and all threats were used in a Markov Chain Monte Carlo generalised linear mixed model (MCMCglmm) to predict IUCN extinction risk category, using phylogenetic and spatial variables as random effects. **b,** Four extinction scenarios were used, baseline; where current, future, and likely to return threats remained as listed by the IUCN^20^, complete abatement; where threats were removed across the entirety of the species range, partial abatement; where threats were removed from at least 50% of the species range, and minimal abatement, where threats were removed from at least 10% of the species range.

**Extended Data Figure 3.**
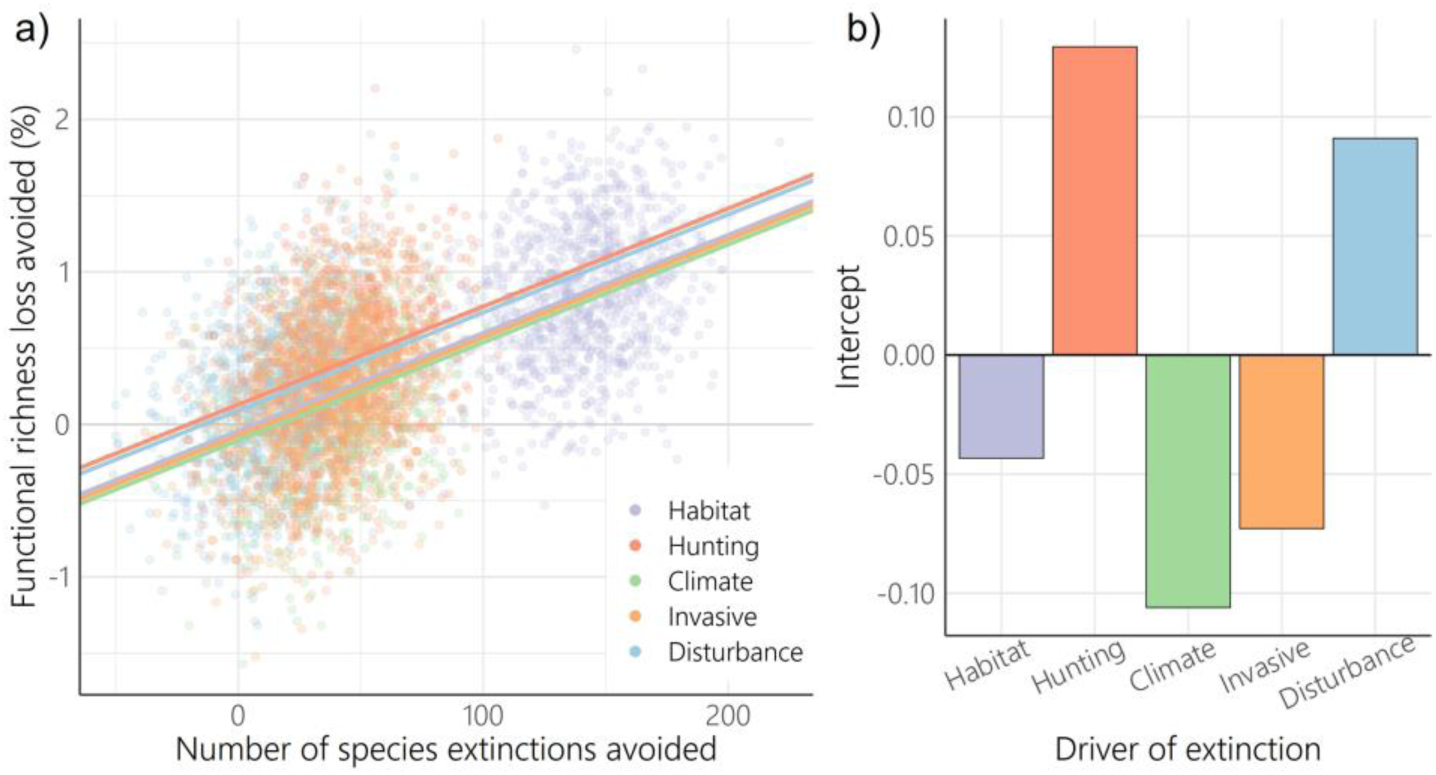
Abatement of hunting and collection, and disturbance and accidental mortality provides disproportionate benefits for functional richness. **a,** Number of species extinctions avoided under driver-specific complete abatement against functional richness loss avoided (% of functional richness of full assemblage) as described by a linear mixed effects model including number of species extinctions avoided and driver of extinction as fixed effects, and iteration number as a random effect. **b,** Intercepts of linear mixed effect model of number of species extinctions avoided against functional richness loss for each driver of extinction showing the proportional impact of each direct driver of extinction given the number of species extinctions. Habitat = habitat loss and degradation, Hunting = hunting and collection, Climate= climate change and severe weather, Invasive = invasive species and disease, Disturbance = disturbance and accidental mortality. Pollution was not included as it made a negligible contribution to functional richness loss (see Extended Data Table 3) (*n* = 5000, 1000 iterations for each extinction scenario).

**Extended Data Figure 4.**
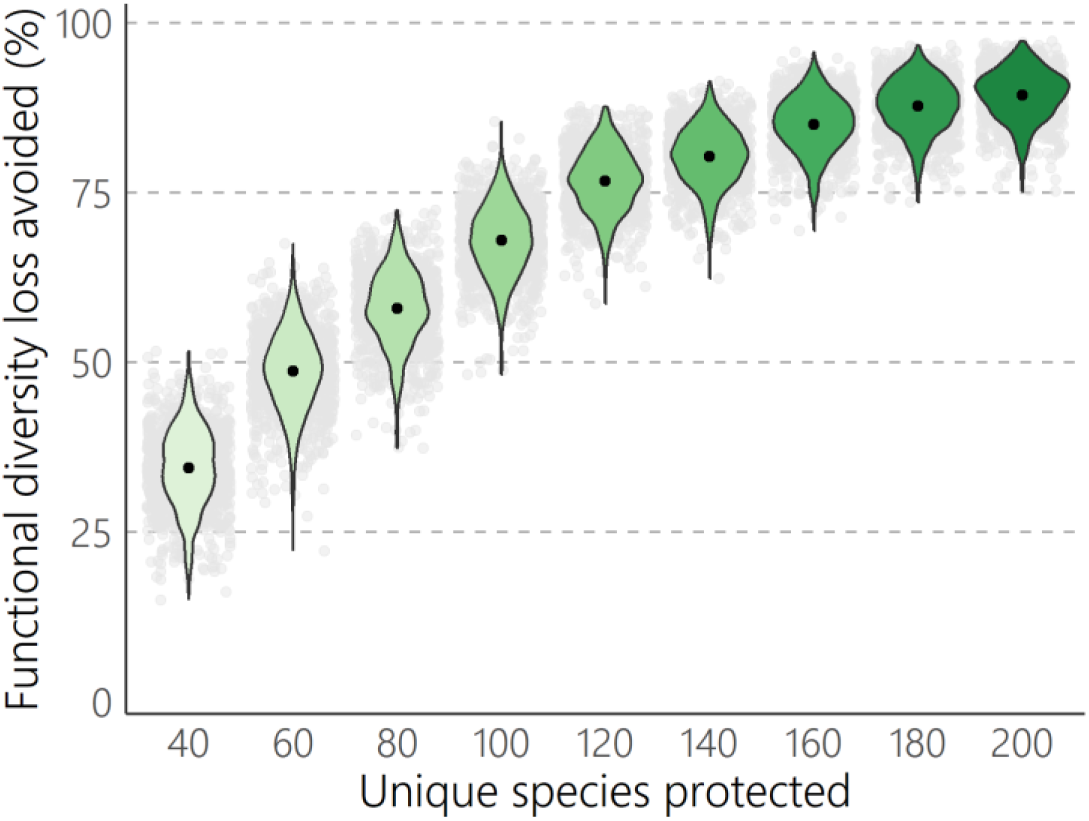
Preventing extinction of unique threatened species reduced projected functional diversity loss. Functional diversity loss avoided (as a percentage of projected functional diversity loss under the baseline scenario) from 1000 iterations. Black points show mean loss avoided; violin plots show the distribution, with grey points showing individual values of loss avoided under each iteration. The number of unique threatened species which were prevented from going extinct (‘protected’) was varied between 40 and 200 unique threatened species at intervals of 20 species.

**Extended Data Figure 5.**
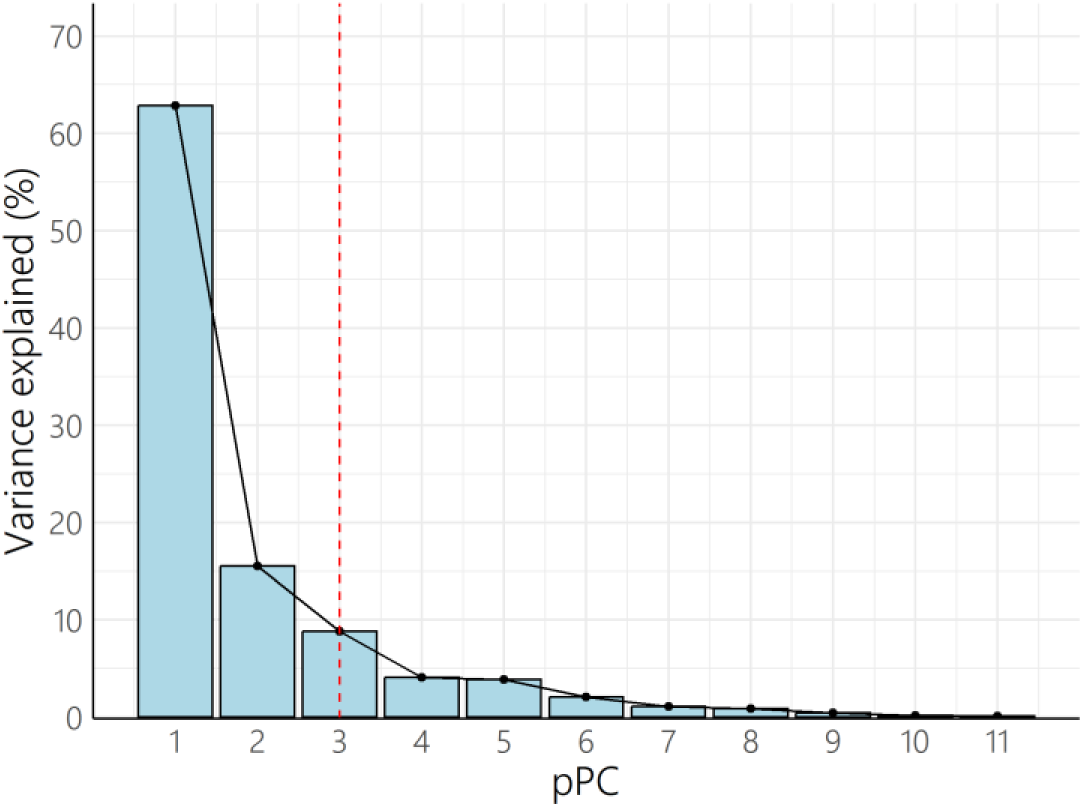
Variance explained by phylogenetic principal components (pPC). Red dotted line indicates elbow after which adding additional phylogenetic principal components would explain little additional variance (*n*=9873 species).

**Extended Data Figure 6.**
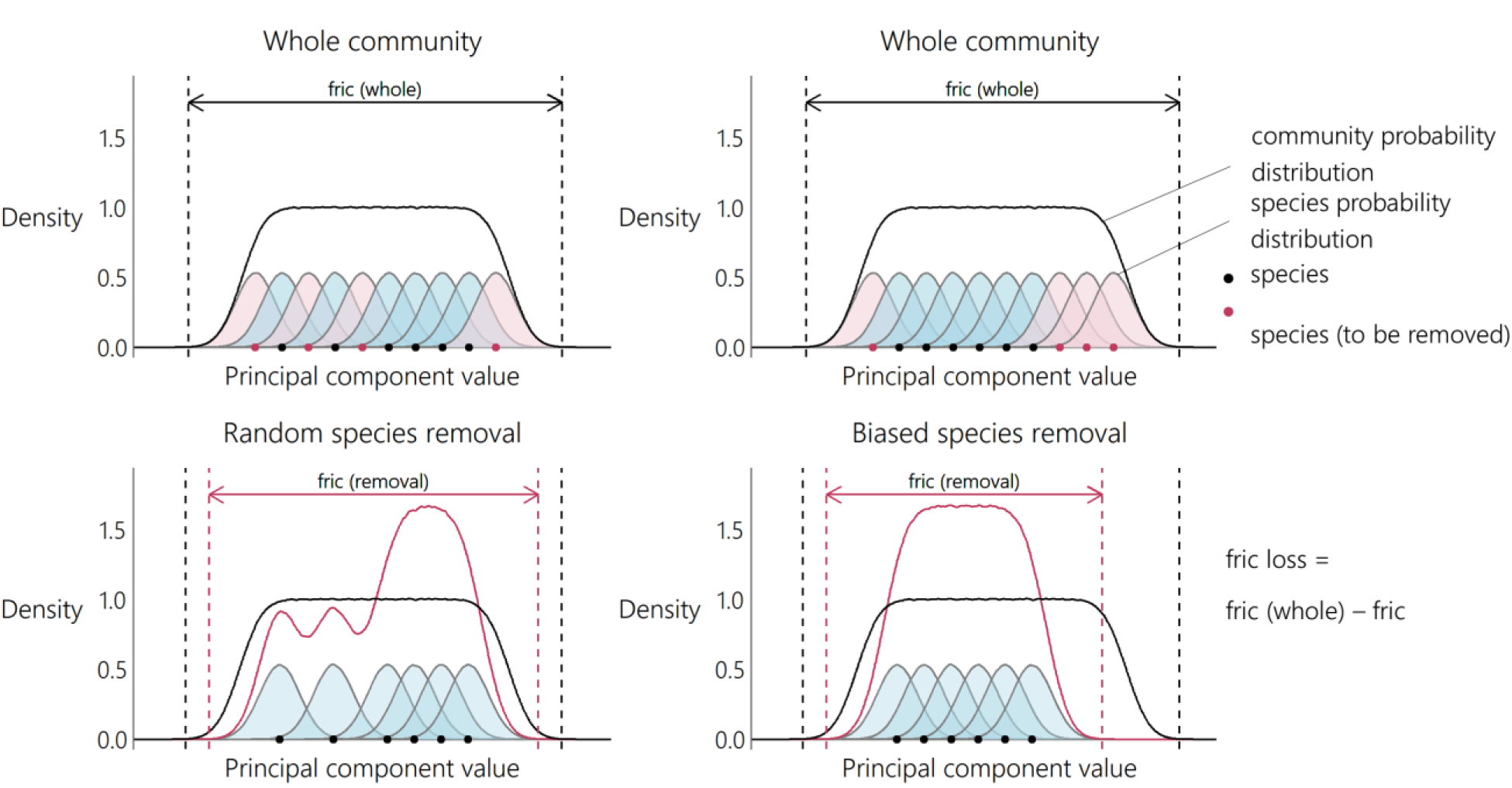
Estimating functional richness loss. When species removal is biased with respect to species traits (principal component values) functional richness loss is greater. Shown in one dimension for simplicity, functional richness was calculated in three-dimensional trait space composed of the first three phylogenetic principal components.

**Extended Data Figure 7.**
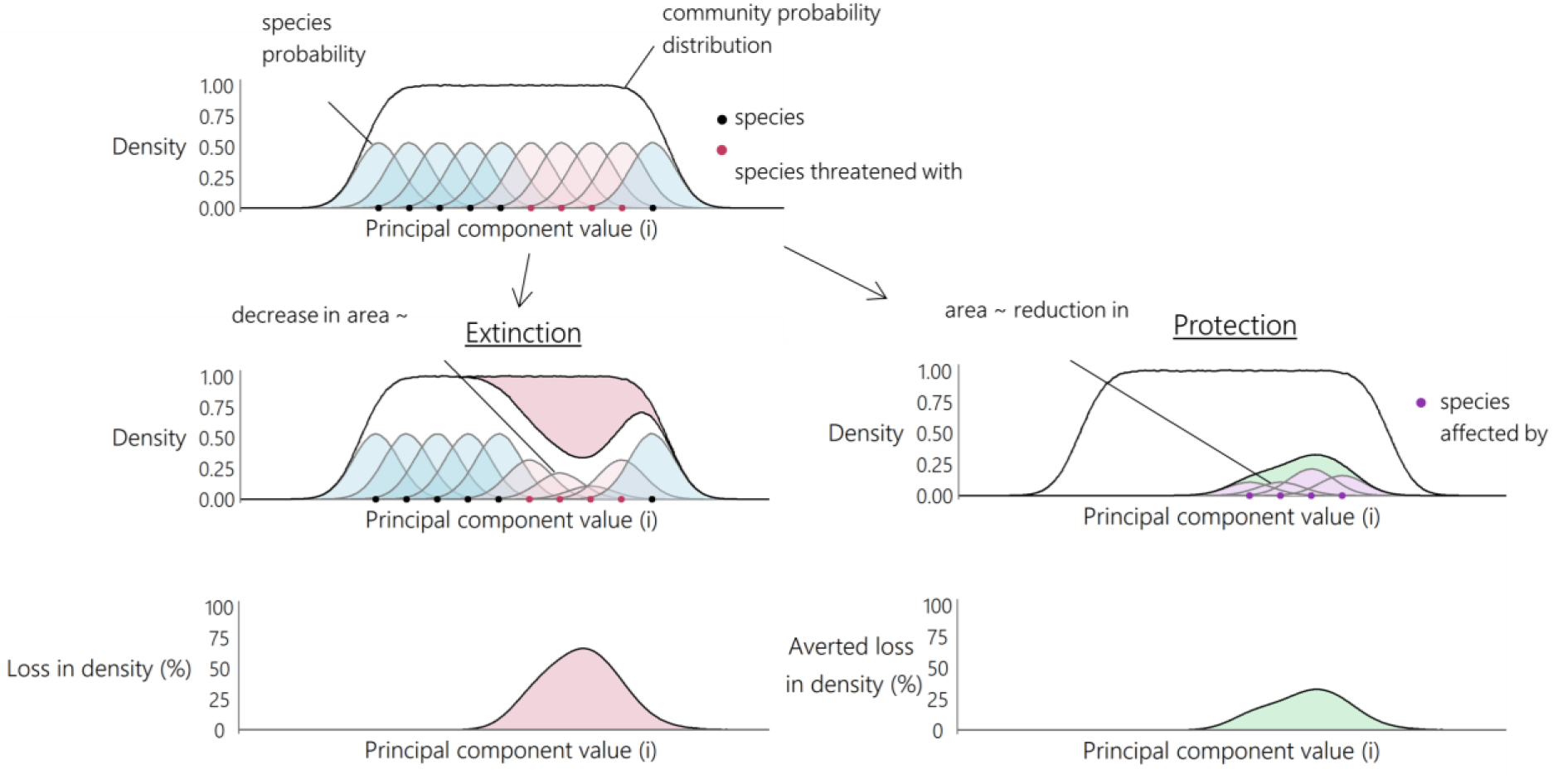
Estimating change in density in morphospace under extinction and conservation. Loss in density of morphospace was calculated using the baseline extinction scenario, and averted loss in density of morphospace was calculated using the complete abatement scenario.

**Extended Data Figure 8.**
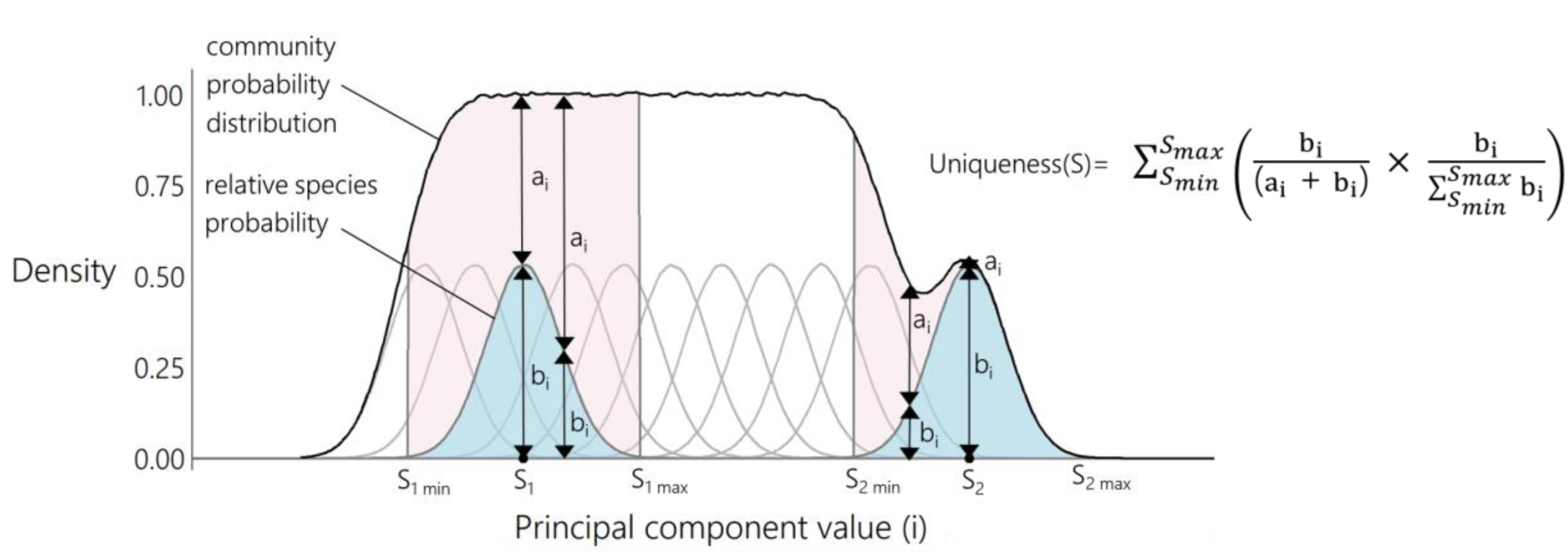
Functional uniqueness calculation. Functional uniqueness describes the proportion of the community probability distribution that was composed of the species probability distribution. Proportions were summed across cells in which the species probability distribution was greater than 0, with a weight proportional to the probability of species occurrence in that cell, indicated by the height of the species probability distribution.

## Notes

### Competing Interest Statement

The authors have declared no competing interest.

