## Supplementary Information for "Threat reduction must be coupled with targeted recovery programmes to conserve global bird diversity"

1. Supplementary Analyses
    a. Extinction risk model description
    b. Predicting downgrading of species extinction risk to Least Concern
    c. Removing the first phylogenetic principal component
    d. Estimating functional diversity with principal components of beak morphology
    e. Reconciling BirdLife and BirdTree taxonomies and including all BirdLife synonyms
    f. Dimensionality and ordination method used in functional richness calculation
    g. Scope and severity imputation accuracy
    h. Prior specification
    i. The impact of past threats that are unlikely to return
2. Supplementary Tables (1-4)
3. Supplementary Figures (1-14)
4. Supplementary References

Supplementary Datasets 1-2 are provided as separate files. Supplementary Datasets and code used for figures and analyses are available at https://figshare.com/s/855f64d2608140f295ab and will be made publicly available upon acceptance.

**Supplementary Dataset 1: Fixed effects of extinction risk model**
Threats to bird species and their significance as fixed effects when explaining variation in extinction risk category using an MCMCglmm. Posterior estimates are given for significant fixed effects, and these threats are grouped into seven threat groups: habitat loss and degradation, hunting and collection, climate change and severe weather, invasive species and disease, disturbance and accidental mortality, pollution and other.

**Supplementary Dataset 2: Top 200 most unique threatened species**
The top 200 most unique threatened bird species ordered by uniqueness, with the scientific name according to Birdlife (Handbook of the Birds of the World and BirdLife

International, 2018), the scientific name according to BirdTree (Jetz et al., 2012), calculated uniqueness values, the IUCN extinction risk category as listed in version 2022-2 (IUCN, 2022) and the main common name listed by the IUCN (IUCN, 2022).

### Supplementary Analyses

##### Extinction risk model description

Four of the five random effects were retained in the extinction risk model. The posterior variance of central latitude was low (mean= 0.934) so was removed from the model, while phylogeny (mean=8.61), centroid longitude (mean=7.89), minimum latitude (mean=7.72), and maximum latitude (mean=6.75) had high posterior variance so were retained in the extinction risk model (Supplementary Fig. 1). Expected population decline from 19 threats (listed by the IUCN second level classification) were useful for explaining variation in extinction risk and retained as fixed effects in the model (Supplementary Dataset 1). Across 1000 iterations the extinction risk category that had the highest posterior probability in the greatest number of iterations matched the correct IUCN category for 86.8% of species. The category that had the highest posterior probability in the greatest or second greatest number of iterations was correct for 99.5% of species. The mean accuracy for a single iteration was 66.0% (standard deviation: 3.5%).

Critically Endangered and Endangered species had a lower accuracy than Near Threatened and Vulnerable species when extinction risk category was assigned using the most frequent category with highest posterior probability (Near Threatened: 98.2%, Vulnerable: 96.9%, Endangered: 59.2%, Critically Endangered: 50.3%). Accuracy when considering the first or second most frequent category with the highest posterior probability was very high for all categories (Near Threatened: 100.0%, Vulnerable: 99.9%, Endangered: 98.9%, Critically Endangered: 97.1%). When extinctions over the next 100 years were predicted according to probabilities assigned to each extinction risk category, the extinction risk model did not over or under predict the number of extinctions relative to those predicted from extinction risk categories listed by the IUCN and using extinction probabilities from Cooke et al. (2019) (model: mean=502 sp., standard deviation=19 sp., IUCN category: 500 sp., standard deviation=12 sp., out of species included in the model). Applying a similar model but with presence/absence data on threats, rather than expected population decline gave slightly lower overall accuracy (86.3%).

**Predicting downgrading of species extinction risk to Least Concern**

Threats contributing to species extinction risk have been comprehensively assessed for all Near Threatened and threatened (including Critically Endangered, Endangered, and Vulnerable) bird species (Mair et al., 2021), but not for Least Concern species. As such the absence of threats for Least Concern species could indicate a genuine absence of threat impact, or an artefact of data recording practices. Least Concern species were not included in the extinction risk model, meaning that the lowest extinction risk the model could predict was Near Threatened. This could lead to an underestimation of the impact of alleviating the drivers of extinction.

To test this, we set the assigned probability ($ex100_c$ in equation 1), of the category Near Threatened to the same as Least Concern (0.0001) for species included in the model and reran the analyses. We found that changing the assigned probability of Near Threatened to match the assigned probability of Least Concern had a small impact on the results. If species were protected from the drivers of extinction across at least half of their range, we predicted that 51% of species extinctions (standard deviation: 6%) (compared to 50% as presented in the main text), and 48% of functional richness loss (standard deviation: 25%, compared to 49% as presented in the main text) could be averted. 377 extinctions (standard deviation: 19 species) and 2.3% decline (standard deviation: 0.3%) in global functional richness remained (as opposed to 385 extinctions and 2.4% decline in functional richness as presented in the main text), with 140 extinctions and 0.9% decline in functional richness directly attributable to the drivers of extinction (compared to 139 extinctions and 0.9% decline in functional richness in the main text). The relative contribution (order of maximum avoidable contributions) that each driver of extinction made to species extinctions and functional richness loss also remain the same (Supplementary Table 1).

Under complete abatement (removal of past threats that were likely to return, ongoing threats, and future threats), 241 species extinctions (standard deviation: 18 species) and 1.4% functional diversity loss (standard deviation: 0.3%) remained (as opposed to 254 species extinctions and 1.5% functional diversity loss when the assigned probability for Near Threatened was 0.01).

**Removing the first phylogenetic principal component**

The first phylogenetic principal component (pPC) explained a large proportion of morphological variation (66%) and was largely a descriptor of body size, explaining variation in wing length, beak length (nares and culmen), beak depth, beak width, body mass, tail length and to some extent tarsus length and Kipp's distance (Extended Data Fig. 1). Body size is correlated with extinction risk in birds (Gaston and Blackburn, 1995) so we removed the first pPC and repeated the analysis, to see if the second and third pPC provided additional information on the impact of human activity on avian diversity.

When only the second and third pPCs were used, we projected that 1.9% of functional diversity will be lost in the next 100 years under current threat levels (standard deviation 1.0%). Protecting species across at least half of their range gave a reduction of 0.6% functional diversity loss (standard deviation: 1.2%, Cohen's D between 50% protection and current threat levels: 0.67), but 1.3% functional diversity loss remained (standard deviation: 0.8%), of which 0.6% functional richness loss was attributable to drivers of extinction (Cohen's D between partial abatement and complete abatement: 1.29). These findings are consistent with the conclusion that protecting species across at least half of their range is effective at alleviating some, but not all of projected functional diversity loss. Drivers of extinction are expected to result in decline in trait variation beyond decline in body size (pPC1) variation, even when half of species ranges are protected from the impact of these threats.

Invasive species and disease, rather than habitat loss and degradation, made the greatest contribution to functional diversity loss when pPC1 was removed (Supplementary Fig. 2). Invasive species and disease resulted in a particularly high loss of functional diversity per number of species extinctions, relative to other threats when functional diversity was measured with only pPC2 and pPC3. While hunting and collection was expected to be particularly biased with respect to body size (pPC1), functional diversity loss attributable to hunting and collection remained particularly high proportional to the number of species extinctions, even when functional diversity was measured with only pPC2 and pPC3 (Supplementary Fig. 3). The contribution of habitat loss and degradation to functional diversity loss was lower when pPC1 was removed, as was the contribution of disturbance and accidental mortality (Supplementary Fig. 3).

**Estimating functional diversity with principal components of beak morphology**

To find if our conclusions were sensitive to the traits used and ways of measuring them, we repeated the analyses on another trait dataset. We tested whether functional diversity loss was prevented by protecting half of species ranges when functional diversity was estimated using principal components of beak morphology measured using three-dimensional scans from Chira et al. (2020), rather than linear measurements of body mass and tail, tarsus, wing, and beak from Avonet (Tobias et al., 2022). While four phylogenetic principal components were available in data obtained from Chira et al. (2020), only the first three principal components were used due to the computation time required to estimate functional diversity in four dimensions, and to enable comparison with the main text.

When functional diversity was measured with beak morphology, we projected that 3.0% of functional diversity will be lost in the next 100 years under current threat levels (standard deviation 0.4%). Protecting species across at least half of their range gave a reduction of 0.8% functional diversity loss (standard deviation: 0.6%, Cohen's D between 50% protection and current threat levels: 2.16), but 1.3% functional diversity loss remained (standard deviation: 0.3%), of which 0.9% functional richness loss was attributable to drivers of extinction (Cohen's D between 50% protection and maximum avoidable contribution of threats: 4.41). These findings are consistent with the conclusion that protecting species across at least half of their range is effective at alleviating some, but not all of projected functional diversity loss.

Habitat loss and degradation made the greatest contribution to functional diversity loss when functional diversity was measured with beak traits (Supplementary Fig. 4), and hunting and collection made the highest contribution proportional to the number of species extinctions (Supplementary Fig. 5), as found in the main text. Disturbance and accidental mortality did not make a high contribution relative to the number of species extinctions when functional diversity was measured with beak traits alone, as observed when using linear measurements tail, tarsus, wing and beak.

#### Reconciling BirdLife and BirdTree taxonomies and including all BirdLife synonyms

When reconciling taxonomic differences between BirdLife (Handbook of the Birds of the World and BirdLife International, 2018) and BirdTree (Jetz et al., 2012), 8958 species had a one-to-one match between BirdLife and BirdTree. A further 878 BirdLife and BirdTree synonym pairs had an exact species name match. 33 synonym pairs were matched which had the same species name but different suffixes (for example *Psittacara holochlorus* [BirdLife] was matched to *Aratinga holochlora* [BirdTree]) or a spelling difference of one letter (*Arses telescopthalmus* [BirdLife] was matched to *Arses telescophthalmus* [BirdTree]). After synonym pairs were matched and removed, 10 one-to-one synonym matches remained and one many-to-one synonym group. For the remaining many-to-one synonym group, one synonym (*Hemignathus affinis*) was selected at random from *Hemignathus affinis* and *Hemignathus hanapepe* (BirdLife) to match with *Hemignathus lucidus* (BirdTree). 24 newly described species in the BirdLife taxonomy were removed as they had no synonyms in the BirdTree taxonomy. This gave 9879 selected synonym matches between BirdLife and BirdTree (89.9% of BirdLife synonyms and 98.9% of BirdTree synonyms). Five species treated as Extinct in the Wild and one species listed as Extinct by IUCN (IUCN, 2022) but not listed as Extinct in the AVONET crosswalk (*Zosterops conspicillatus*) were removed from the analysis giving a total of 9873 species which were included in the main analyses.

Phylogenetic principal component analysis was used to summarise trait variation whilst accounting for covariance between species. To carry out phylogenetic principal component analysis and simulate extinction scenarios we required a one-to-one match between synonyms in the BirdLife taxonomy (used by the IUCN Red List) and the BirdTree taxonomy (used by the Jetz taxonomy, Jetz et al., 2012). We were able to assign one-to-one matches for 9873 species, which represents 89.9% of synonyms under the BirdLife taxonomy and 98.9% of synonyms under the BirdTree taxonomy. It was not possible to include multiple BirdLife data points per tip when carrying out phylogenetic principal component analysis due to pseudo-replication.

To test the impact of not including all BirdLife synonyms we relaxed the requirement that all principal components should be strictly orthogonal when accounting for phylogenetic covariance, and conducted principal component analysis with all BirdLife synonyms (using the function *prcomp*), but not accounting for phylogenetic covariance between species. We also carried out principal component analysis without accounting for phylogenetic covariance for the 9873 species included in the main text. We compared projected species richness loss and functional richness loss when using 9873 species, and the 10 993 extant species as listed under the BirdLife taxonomy. We constructed an extinction risk model (MCMCglmm) using all BirdLife synonyms and all BirdTree synonyms (Jetz et al., 2012). For BirdLife synonyms with multiple tips, we took the mean extinction probability across the tips. We simulated 1000 iterations of the baseline and management scenarios for all drivers of extinction.

Next, we compared projected functional richness loss when using principal component analysis without accounting for phylogenetic covariance, and when using a phylogenetic principal component analysis, with 9783 species. We predicted extinctions under the baseline scenario, with complete abatement of all drivers of extinction, and partial abatement (50%) of all drivers of extinction (with 1000 iterations for each scenario) and quantified their impact on functional richness (mean number of projected extinctions should not change). We compared the results to those calculated using phylogenetic principal component analysis to assess the relative impact of not using phylogenetic principal component analysis and not including all BirdLife species.

When using all species, we projected 5.5 ± 0.2% loss in species richness (mean of 605 species extinctions) in the next 100 years under the baseline scenario (standard deviation: 20 sp.). As a percentage of species included, species richness loss was slightly higher when using all species compared to when only 9873 species were included (all species: mean -5.5%, standard deviation 0.2%, 9873 species: mean -5.3%, standard deviation 0.2%). Under complete abatement with all species 2.8% of species were projected to go extinct (standard deviation: 0.2%), compared to 2.6% when 9873 species were included (standard deviation: 0.2%). Similarly, projected extinctions under partial abatement were slightly higher when using all species rather than 9873 species (-4.1% and -3.9% respectively, standard deviation 0.2% for both). Functional richness loss was slightly greater in the baseline scenario and under complete abatement when using all synonyms, but was always within one standard deviation of the mean functional richness loss when using 9873 species. There was negligible difference in the partial abatement scenario (-1.3% for both, standard deviation 0.3%). These results show that including only species for which we could assign a one-to-one match between BirdLife and BirdTree had a small impact on results and did not affect our conclusions.

Using traditional principal component analysis rather than phylogenetic principal component analysis had a small impact on projected functional richness loss under a baseline scenario (Supplementary Fig. 6). With phylogenetic principal component analysis, we projected functional richness to change by -3.2% compared to -3.1% with a non-phylogenetic (traditional) principal component analysis (standard deviation 0.4% for both). Under complete abatement the projected functional richness change was -1.5% for analysis with phylogenetic principal component analysis and with -1.3% non-phylogenetic principal component analysis (standard deviation: 0.3% for both). Under partial abatement the projected functional diversity change was -2.3% for analysis with phylogenetic principal component analysis and -2.2% for analysis with non-phylogenetic principal component analysis (standard deviation: 0.3% and 0.4% respectively).

**Dimensionality and ordination method used in functional richness calculation**

We conducted sensitivity analysis to test the effect of using a different number of dimensions (principal components) and different ordination methods on functional richness estimations.

We calculated functional richness in the baseline, complete abatement, partial abatement and driver-specific abatement scenarios using two and four principal components and compared the results to those obtained using three principal components (as presented in the main text). Calculating functional richness with ~10000 species in four dimensions is very computationally demanding so it was necessary to reduce the number of divisions that each dimension (principal component) was divided into when estimating trait probability densities (specified using the *n_divisions* parameter in the function *TPDsMean* in R package *TPD* [Carmona, 2019]). When using four dimensions we used 25 divisions per principal component and compared to functional richness estimations calculated using three dimensions and 25 divisions. When using two dimensions we used 50 divisions and compared to functional richness estimations calculated using three dimensions and 50 divisions (as in the main text).

All extinction scenarios were run 100 times rather than 1000 times as in the main text due to computational limitations. We compared functional richness loss under baseline, partial and complete abatement scenarios between different dimensions and divisions using a three-way Analysis of Variance. To find the severity of functional richness loss under a given driver of extinction in relation to the number of species projected to go extinct, we used a linear mixed effects model to describe the functional richness loss avoided, using the number of species extinctions avoided, and the driver of extinction as predictor variables, and model iteration as a random effect (as described in the main text), with a separate model for each combination of dimensions and divisions used.

Using a greater number of dimensions and a greater number of divisions increased projected diversity loss ($p<0.001$ for both variables, $n=1200$), but the variance explained by the number of dimensions ($SS=188.1$, $df=2$) and divisions ($SS=71.7$, $df=2$), was low relative to that described by the extinction scenario ($SS=629.8$, $df=2$). Projected functional richness loss under the baseline scenario ranged from 2.4 ± 0.9% with two dimensions and 50 divisions, to 3.8 ± 0.4% with four dimensions and 25 divisions. The proportion of projected functional diversity loss avoided varied between 54.0 ± 10.3% and 64.6 ±21.2% under complete abatement, and between 20.2 ± 56.3% and 34.9 ± 20.3% under partial abatement (Supplementary Table 2, Supplementary Fig. 7a). Habitat loss and degradation and hunting and collection resulted in the greatest functional diversity loss regardless of the number of divisions or number of dimensions used, although hunting and collection resulted in greater projected functional diversity loss than habitat loss and degradation when functional richness was estimated with two dimensions and 50 divisions (Supplementary Fig. 7a). Hunting and collection had a disproportionate impact on functional richness relative to the number of species extinctions regardless of the number of dimensions and divisions used ($p<0.001$, $n=500$), and disturbance and accidental mortality had a disproportionate impact on functional richness when using three or four dimensions and 50 divisions, but not when using two or three dimensions and 25 divisions ($p>0.05$, $n=500$). While the number of dimensions and the number of divisions did affect estimations of projected functional richness loss, our conclusions that partial abatement will only be partially effective at averting projected diversity loss, and that abatement of habitat loss and degradation and hunting and collection will avoid the greatest proportion of projected diversity loss relative to other drivers, remain.

We compared projected functional richness loss when trait space was constructed using principal components analysis (PCA, as presented in the main text), principal coordinates analysis, and non-metric dimensional scaling. We conducted principal coordinates analysis (PCoA) based on Manhattan distances. We used Manhattan distances rather than Euclidean distances as PCoA and principal components analysis (PCA) are identical when principal components are estimated based on covariance and principal coordinates are estimated based on Euclidean distances. We conducted PCoA using the *pcoa* function in the package *ape* (Paradis and Schliep, 2019) using Lingoes procedure (Lingoes, 1971) to correct for negative eigenvalues. Three principal coordinates were optimal for describing variation in Manhattan distances between species (identified using a scree plot through the package *pathviewr* [Baliga et al., 2021], Supplementary Fig. 8).

We conducted NMDS using *metaMDS* function in the package *vegan* (Oksanen et al., 2024) using the *monoMDS* engine, based on Manhattan distance matrix with 40 starts and 3 dimensions. The function was not able to obtain the same best result twice with different starts (even with multiple runs and increasing to 100 starts) indicating that the obtained ordination could represent a local rather than global optima. Despite this, the difference in stress between starts was low (0.05) so it is likely that ordinations with different starts were similar, and ordination adequately represented the original data.

All extinction scenarios were run 100 times. We compared functional richness loss under baseline, partial and complete abatement scenarios between different ordination methods using a two-way Analysis of Variance. To find the severity of functional richness loss under a given driver of extinction in relation to the number of species projected to go extinct, we used a linear mixed effects model as described above, with a separate model for each ordination method.

While PCoA and NMDS resulted in slightly lower projected functional richness loss under the baseline, complete abatement and partial abatement scenarios than PCA, the ordination method explained a low proportion of variance compared to the extinction scenario (ordination method SS: 51.9, df: 2, extinction scenario SS: 357.5, df: 2) and the proportion of projected functional richness loss avoided was similar between methods (Supplementary Fig. 9a, Supplementary Table 3). Habitat loss and degradation and hunting and collection resulted in the greatest functional diversity loss regardless of the ordination method used (Supplementary Fig. 9b). Hunting and collection had a disproportionate impact on functional richness relative to the number of species extinctions regardless of the ordination method used ($p<0.001$, $n=500$), and disturbance and accidental mortality had a disproportionate impact on functional richness when using PCA or NMDS, but not when using PCoA ($p>0.05$, $n=500$). As with the number of dimensions and number of divisions, ordination method did not affect our conclusions that partial abatement will only be partially effective at averting projected diversity loss, and that abatement of habitat loss and degradation and hunting and collection will avoid the greatest proportion of projected diversity loss relative to other drivers.

**Scope and severity imputation accuracy**

Across all species, 11.58% of threat data (threat-species combinations) were missing scope or severity values. Missing scope and severity data were imputed with missForest imputation (implemented through R package *missForest* [Stekhoven, 2022]) from threat type, scope, severity and timing, and phylogenetic eigenvectors (Debastiani et al., 2021). To test the accuracy of imputation, scope and severity data were removed from the complete dataset (a subset of the master dataset including only species and threat combinations where all of scope, severity and timing were present), in a manner that mimicked the structure of missingness in the master dataset.

In the master dataset 0.12% of threat-species combinations were missing data across all of scope, severity and timing, so all of scope, severity and timing were removed for this proportion in the complete dataset. Likewise, 2.04% of threat-species combinations were missing both scope and severity in the master data, so 2.04% of species-threat combinations had scope and severity removed in the complete dataset. Following this, 11.19% severity data were removed at random across threat-species combinations to mimic the proportion of data missing in the master dataset and 2.48% of scope data were removed. Removal of data and imputation was repeated 100 times. The percentage of true values in the complete dataset that matched imputed values from each removal dataset was calculated.

Overall, the mean accuracy of imputation across 100 iterations of data removal was 82.5% (standard deviation: 0.8%). Mean imputation accuracy was slightly higher for severity (mean: 82.8%, standard deviation: 0.8%) than scope (mean: 81.5%, standard deviation: 1.7%). When two of scope, severity or timing were missing, mean imputation accuracy was 80.0% (standard deviation 1.7%), and when all three of scope, severity and timing were missing, mean imputation was 75.2% (standard deviation

7%). Due to high imputation accuracy we chose to proceed with imputing missing scope, severity and timing values.

##### Prior specification

We tested the sensitivity of the extinction risk model to random effect prior specification. Firstly, we tested the impact of using a parameter expanded prior instead of a Chi-squared prior for the phylogenetic random effect. Cauchy-scaled priors were used for fixed effects as in the main text, the residual variance was fixed at 1, and parameter expanded priors used for spatial variables). The model was run for 103 000 iterations and convergence was checked using plots of parameter traces. The posterior distribution of phylogenetic variance was similar whether a Chi-squared or parameter expanded prior were used (Supplementary Fig. 10), as were posterior distributions of fixed effects (Supplementary Fig. 11). Projected extinctions under the baseline (Chi-squared prior: 517 ± 19 species, parameter expanded prior: 517 ± 18 species), complete abatement (Chi-squared: 254 ± 19 species, parameter expanded prior: 254 ± 18 species) and partial abatement (Chi-squared prior: 385 ± 18 species, parameter expanded prior: 385 ± 18 species) scenarios were similar when using a model with a Chi-squared or parameter expanded prior for the phylogenetic random effect.

We then tested the impact of using inverse-Wishart priors with an expected covariance of 1 and a degree of belief of 0 for the spatial random effects rather than parameter expanded priors. Cauchy-scaled priors were used for fixed effects, as in the main text, the residual variance was fixed at 1, and a Chi-squared prior was used for the phylogenetic random effect. The model was run for 103 000 iterations and convergence was checked using plots of parameter traces. As expected, when using the Inverse-Wishart prior the estimated variance was closer to zero for spatial random effects, than when using the parameter expanded prior (Supplementary Fig. 12) (Hadfield, 2017) but posterior distributions of fixed effects were similar regardless of random effect prior specification (Supplementary Fig. 13). Projected extinctions under the baseline (parameter expanded prior: 517 ± 19 species, inverse-Wishart prior: 518 ± 18 species), complete abatement (parameter expanded prior: 254 ± 19 species, inverse-Wishart prior: 252 ± 18 species) and partial abatement (parameter expanded prior: 385 ± 18 species, inverse-Wishart prior: 384 ± 18 species) scenarios were similar when using a model with a parameter expanded or inverse-Wishart prior for the spatial random effect.

We used weak priors for fixed effects. Posterior distributions of fixed effects had a much narrower variance than prior distributions, and the mean of posterior distributions often differed from that of the prior distributions (mean=0) (Supplementary Fig. 14), indicating that fixed effect priors did not unduly influence posterior estimates.

**The impact of past threats that are unlikely to return**

The precise impact of past threats that are unlikely to return on species extinction risk is unknown. As most threats only affected a small number of species under the timing "*Past, Unlikely to Return*" it was not possible to estimate the individual effect of each threat in the past on extinction risk separately to their current and future impact. As such, past threats that were unlikely to return were grouped with current and future threats when modelling extinction risk. In the main text, we assumed that past threats that were unlikely to return were still important for explaining the extinction risk of a species. When abatement scenarios were applied, we did not remove the impact of threats with a timing of "*Past, Unlikely to Return*" so that the impact of listed past threats remained constant between baseline and abatement extinction scenarios. However, if the impact of past threats can be completely removed, as assumed for current and future threats in the complete abatement scenario, past threats may have no impact on species extinction risk in the present. We tested how removing the impact of past threats from both the extinction risk model, and extinction scenarios affected our results.

We fitted the extinction risk model including 19 IUCN threat categories as fixed effects as described in the main text but excluding the impact of threats with a timing of "*Past, Unlikely to Return*". One phylogenetic and three spatial random effects (including minimum latitude, maximum latitude and centroid longitude) were used, with a chi-squared prior for the phylogenetic random effect, and parameter expanded priors for the spatial random effects. For the fixed effects we used Cauchy-scaled Gelman priors. MCMC chains were run for 103000 iterations, with a burn in period of 3000 iterations and a thinning interval of 100 iterations. We calculated the self-prediction accuracy of the model through calculating the proportion of species for which the category listed by the IUCN (2022) matched the category that most frequently had the highest probability across iterations. Finally, we projected species extinctions and functional richness loss in the next 100 years under the baseline and complete abatement scenario. Contrary to the main text, in this sensitivity analysis we assumed that past threats that were unlikely to return did not contribute to extinction risk, so they were not included when projecting biodiversity loss under either the baseline or complete abatement scenario.

The model without past threats had marginally higher self-prediction accuracy (87.5%) than the model with past threats (as presented in the main text, 86.8%). Nevertheless, the assumption of whether past threats were important or irrelevant for predicting present extinction risk had no impact on projected biodiversity loss under the baseline or complete abatement scenario (Supplementary Table 4).

### Supplementary Tables

**Supplementary Table 1.** Mean number of species extinctions and functional richness loss attributable to each threat (standard deviation in brackets) when Near Threatened was assigned a probability of 0.0001 (equal to Least Concern) and when Near Threatened was assigned a probability of 0.01 (as in main text) for species included in the model.

| | Species extinctions | | Functional richness loss | |
|---|---|---|---|---|
| Driver of extinction | Near Threatened: 0.0001 | Near Threatened: 0.01 | Near Threatened: 0.0001 | Near Threatened: 0.01 |
| Habitat | 149 (24) | 141 (24) | 0.89 (0.47) | 0.86 (0.46) |
| Hunting | 51 (23) | 42 (23) | 0.47 (0.48) | 0.40 (0.48) |
| Climate | 39 (23) | 37 (22) | 0.13 (0.48) | 0.13 (0.46) |
| Invasive | 38 (22) | 37 (22) | 0.19 (0.47) | 0.16 (0.48) |
| Disturbance | 18 (23) | 15 (23) | 0.21 (0.47) | 0.19 (0.49) |
| Pollution | 17 (22) | 8 (22) | 0.07 (0.47) | 0.04 (0.46) |

**Supplementary Table 2**. Projected functional diversity loss avoided under partial and complete abatement (% of projected functional richness loss under baseline scenario, mean ± standard deviation). 100 iterations were run for each extinction scenario.

| Extinction scenario | Two dimensions 50 divisions | Three dimensions 50 divisions | Three dimensions 25 divisions | Four dimensions 25 divisions |
|---|---|---|---|---|
| Partial abatement | 20.2 ± 56.3% | 26.0 ± 12.1% | 34.9 ± 20.3% | 28.1 ± 8.5% |
| Complete abatement | 58.7 ± 34.7% | 54.0 ± 10.3% | 64.6 ±21.2% | 55.4 ± 8.1% |

**Supplementary Table 3**. Projected functional diversity loss avoided under partial and complete abatement (% of projected functional richness loss under baseline scenario, mean ± standard deviation). 100 iterations were run for each extinction scenario.

| Extinction scenario | PCA | PCoA | NMDS |
|---|---|---|---|
| Partial abatement | 26.0 ± 12.1% | 26.1 ± 17.5% | 25.8 ± 13.7% |
| Complete abatement | 54.0 ± 10.3% | 55.0 ± 12.4% | 55.2 ± 12.2% |

**Supplementary Table 4.** Species extinctions and functional richness loss under the baseline scenario and the complete abatement scenario, when past threats are included in the extinction risk model (with past threats) and contribute to extinction risk in extinction scenarios, and when past threats are not included in the extinction risk model and do not contribute to extinction risk in extinction scenarios (without past threats). Complete abatement refers to biodiversity loss remaining after complete abatement (rather than the reduction in biodiversity loss).

| | Species extinctions | | Functional richness loss | |
|---|---|---|---|---|
| Extinction scenario | With past | Without past | With past | Without past |
| Baseline | 5.2 ± 0.2% (517 ± 19 sp.) | 5.2% ± 0.2% (518 ± 17 sp.) | 3.2 ± 0.4% | 3.2 ± 0.4% |
| Complete abatement | 2.6 ± 0.2% (254 ± 19 sp.) | 2.7% ± 0.2% (254 ± 17 sp.) | 1.5 ± 0.3% | 1.5 ± 0.3% |

### Supplementary Figures

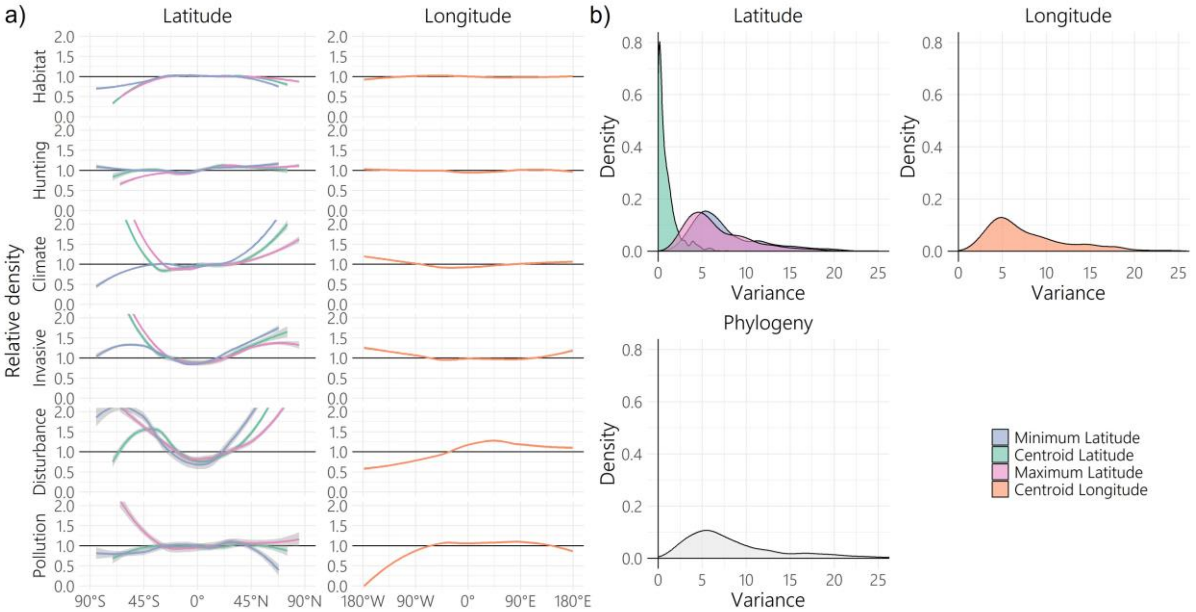


**Supplementary Figure 1. Spatial variables explain variation in extinction risk. a**, Drivers of extinction are unevenly distributed with respect to spatial variables (minimum latitude, maximum latitude and centroid longitude) in Near-Threatened and threatened species. Relative density describes the density distribution of Near-Threatened and threatened species affected by a threat, divided by the density distribution of all Near-Threatened and threatened species included in the study (2087 species). Values above 1 indicate higher prevalence of threat at a given latitude or longitude than expected from the spatial distribution of Near-Threatened and threatened species. **b,** Posterior distribution of five variance components in a model of extinction risk category with threats affecting Near-Threatened and threatened species as fixed effects and with centroid latitude, centroid longitude, maximum latitude, minimum latitude and species (phylogeny) as random effects (referred to as the extinction risk model in the main text, 2087 species, 1000 posterior estimates). Centroid latitude was not included in the final model owing to low posterior variance.

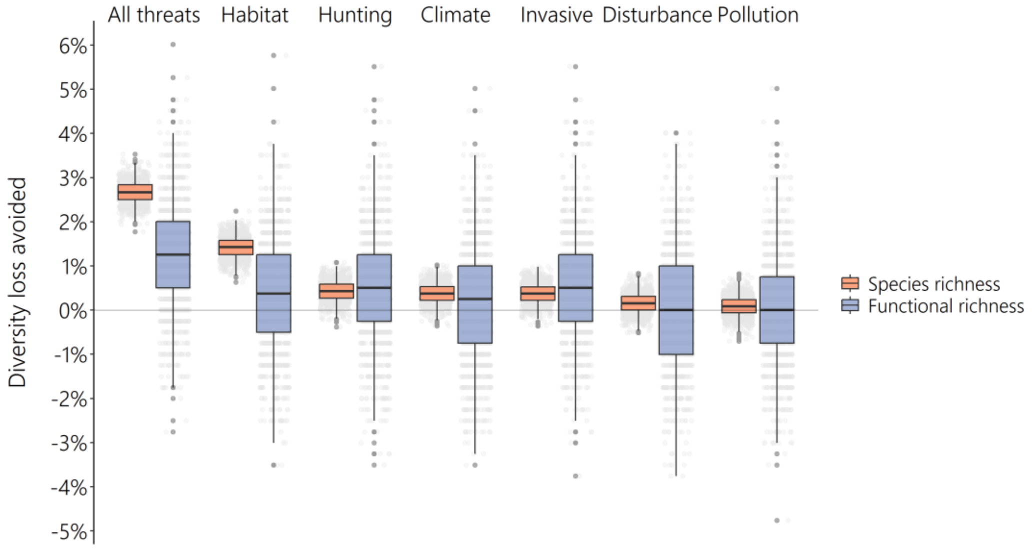


**Supplementary Figure 2. Diversity loss avoided under driver-specific complete abatement of all drivers of extinction when pPC1 was not included.** Maximum avoidable contribution of species richness loss (orange) and functional richness loss (purple) for each driver of extinction when functional richness was measured with pPC2 and pPC3 only. Lower hinges show the 25th percentile and lower whiskers show the minimum value that is within 1.5 times the interquartile range below the 25th percentile. Upper hinges show the 75th percentile and upper whiskers show the maximum value that is within 1.5 times the interquartile range above the 75th percentile. Grey points show diversity loss avoided in each iteration (1000 iterations were run for each extinction scenario). All threats = all threats, including those not included in six major categories, Habitat = habitat loss and degradation, Hunting = hunting and collection, Climate= climate change and severe weather, Invasive = invasive species and disease, Disturbance = disturbance and accidental mortality, Pollution = pollution.

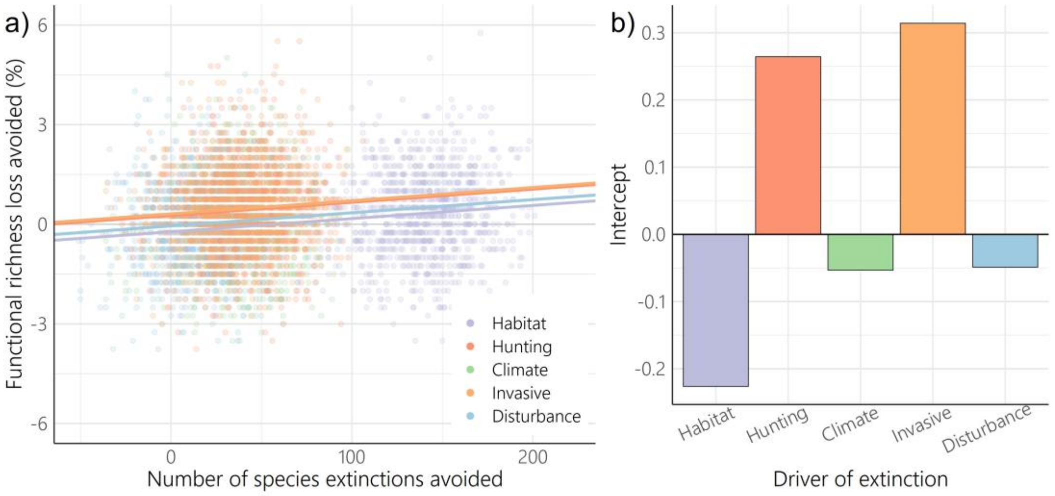


**Supplementary Figure 3. Abatement of hunting and collection and invasive species and disease provides disproportionate benefits for functional richness, when pPC1 was not included.** Functional richness was quantified using pPC2 and pPC3 only. **a,** Number of species extinctions avoided under driver-specific complete abatement against functional richness loss avoided (% of functional richness of full assemblage) as described by a linear mixed effects model including number of species extinctions avoided and driver of extinction as fixed effects, and iteration number as a random effect. **b,** Intercepts of linear mixed effect model of number of species extinctions avoided against functional richness loss for each driver of extinction showing the proportional impact of each direct driver of extinction given the number of species extinctions. Habitat = habitat loss and degradation, Hunting = hunting and collection, Climate= climate change and severe weather, Invasive = invasive species and disease, Disturbance = disturbance and accidental mortality. Pollution was not included as it made a negligible contribution to functional richness loss (see Extended Data Table 3) (*n* = 5000, 1000 iterations for each extinction scenario).

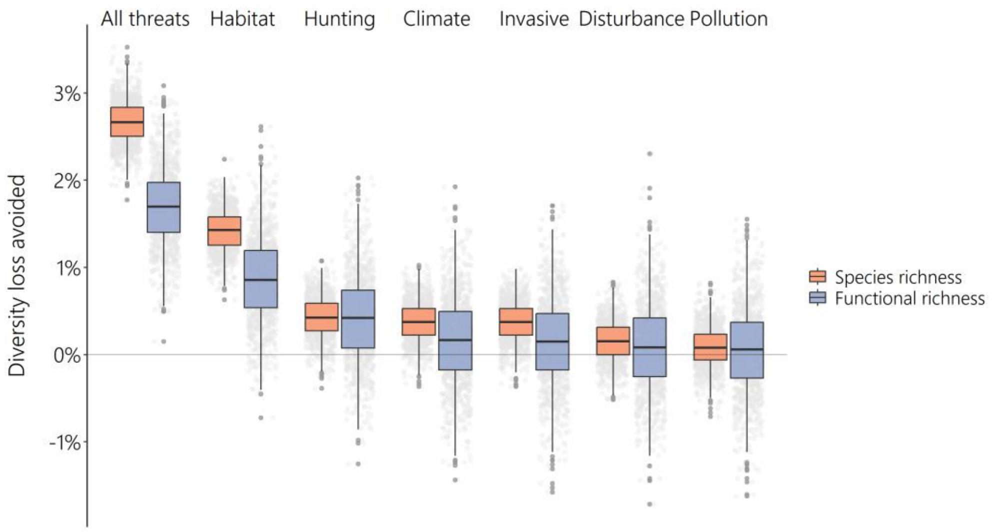


**Supplementary Figure 4. Diversity loss avoided under driver-specific complete abatement of all drivers of extinction when functional richness was estimated from beak shape.** Maximum avoidable contribution of species richness loss (orange) and functional richness loss (purple) for each driver of extinction when functional richness was measured with the first three phylogenetic principal components of beak shape from Chira et al. (2022). Lower hinges show the 25$^{th}$ percentile and lower whiskers show the minimum value that is within 1.5 times the interquartile range below the 25$^{th}$ percentile. Upper hinges show the 75$^{th}$ percentile and upper whiskers show the maximum value that is within 1.5 times the interquartile range above the 75$^{th}$ percentile. Grey points show diversity loss avoided in each iteration (1000 iterations were run for each extinction scenario). All threats = all threats, including those not included in six major categories, Habitat = habitat loss and degradation, Hunting = hunting and collection, Climate= climate change and severe weather, Invasive = invasive species and disease, Disturbance = disturbance and accidental mortality, Pollution = pollution.

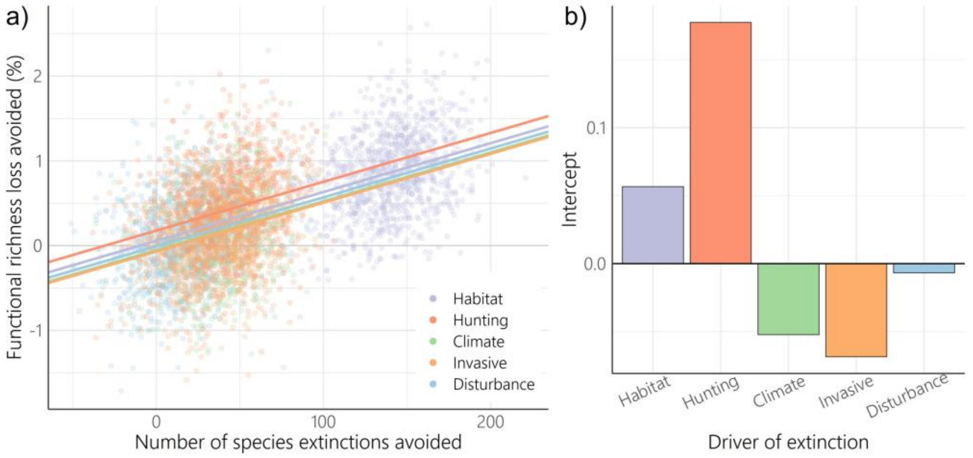


**Supplementary Figure 5. Abatement of hunting and collection provides disproportionate benefits for functional richness, but disturbance and accidental mortality does not, when functional richness was estimated from beak shape.** Functional richness was quantified with the first three phylogenetic principal components of beak shape from Chira et al. (2022). **a,** Number of species extinctions avoided under driver-specific complete abatement against functional richness loss avoided (% of functional richness of full assemblage) as described by a linear mixed effects model including number of species extinctions avoided and driver of extinction as fixed effects, and iteration number as a random effect. **b,** Intercepts of linear mixed effect model of number of species extinctions avoided against functional richness loss for each driver of extinction showing the proportional impact of each direct driver of extinction given the number of species extinctions. Habitat = habitat loss and degradation, Hunting = hunting and collection, Climate= climate change and severe weather, Invasive = invasive species and disease, Disturbance = disturbance and accidental mortality. Pollution was not included as it made a negligible contribution to functional richness loss (see Extended Data Table 3) (*n* = 5000, 1000 iterations for each extinction scenario).

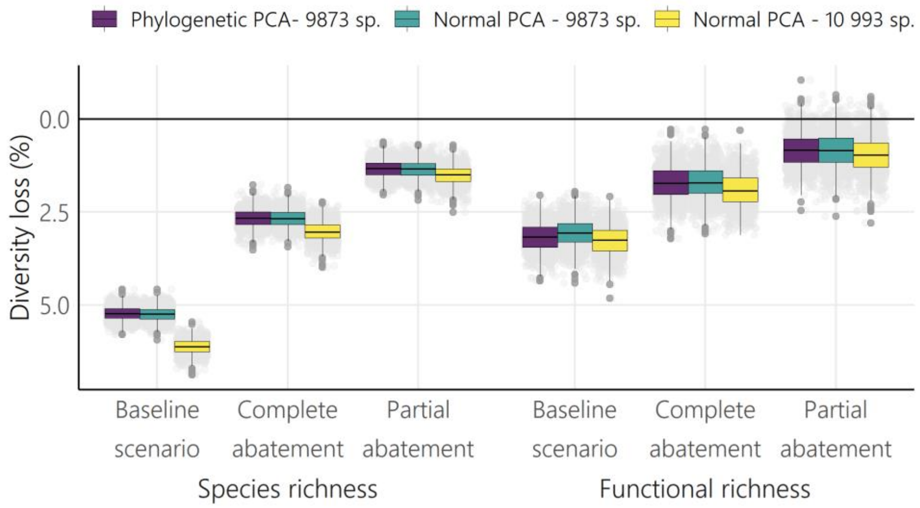


**Supplementary Figure 6. Projected diversity loss when including all synonyms.** Species richness and functional richness loss under the baseline scenario, complete abatement of all drivers of extinction (impact of threats removed across the entirety of species ranges), and partial abatement of all drivers of extinction (impact of threats removed across at least 50% of species ranges). The analysis was run three times, once with a phylogenetic principal component analysis (PCA) including 9873 species that could be matched between the taxonomy used by the Jetz et al. (2012) phylogeny, and the BirdLife taxonomy (used by IUCN), once with the same species but using a traditional PCA (not a phylogenetic PCA) and once with a traditional PCA using all species as listed under the BirdLife taxonomy. Lower hinges show the 25th percentile and lower whiskers show the minimum value that is within 1.5 times the interquartile range below the 25th percentile. Upper hinges show the 75th percentile and upper whiskers show the maximum value that is within 1.5 times the interquartile range above the 75th percentile. Grey points show diversity loss avoided in each iteration (1000 iterations were run for each extinction scenario).

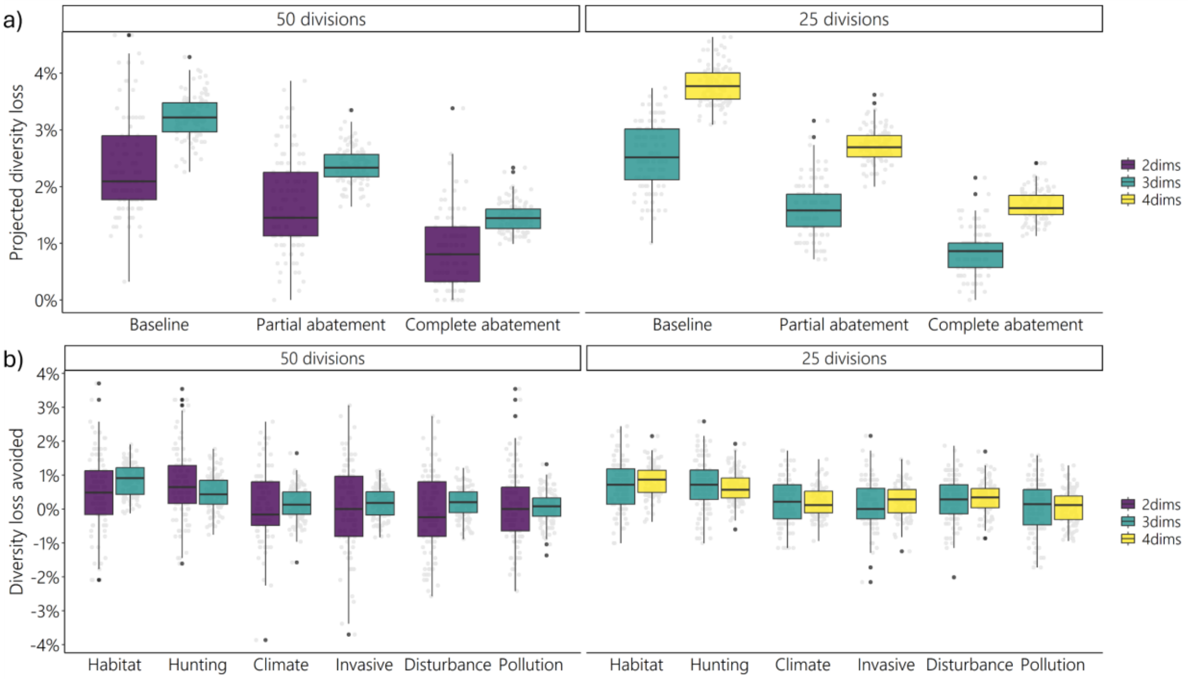


**Supplementary Figure 7. Impact of number of dimensions and divisions on functional richness estimations.** Functional richness was estimated using two dimensions (2dims), three dimensions (3dims) and four dimensions (4dmis), with 50 divisions of principal components for two and three dimensions, and 25 divisions of principal components for three and four dimensions. Grey points show raw data with 100 iterations for each extinction scenario. a) Projected diversity loss under baseline, partial abatement and complete abatement scenarios, and b) Diversity loss avoided under driver-specific abatement. Error bars of diversity loss avoided can go below zero if some iterations of driver-specific abatement have greater projected functional diversity loss than some iterations of baseline extinction scenarios.

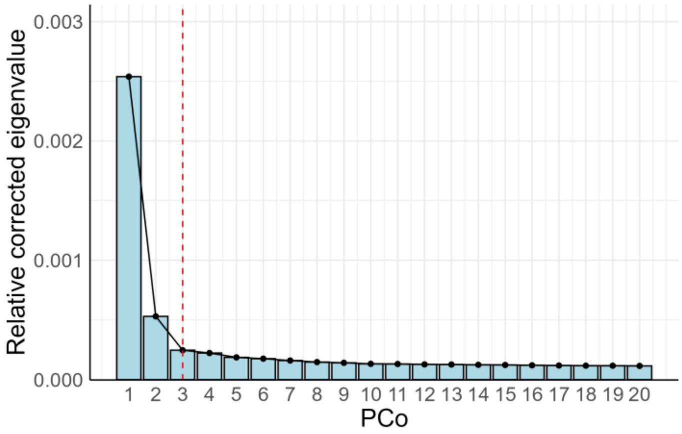


**Supplementary Figure 8. Three principal coordinates (PCo) provide optimal explanation of variation in Manhattan distances between species.** Scree plot showing the relative corrected eigenvalues of the first 20 principal coordinates obtained using principal coordinates analysis. Red dotted line indicates elbow after which adding additional phylogenetic principal components would explain little additional variance (n=9873 species). Eigenvalues were corrected using Lingoes procedure (Lingoes, 1971).

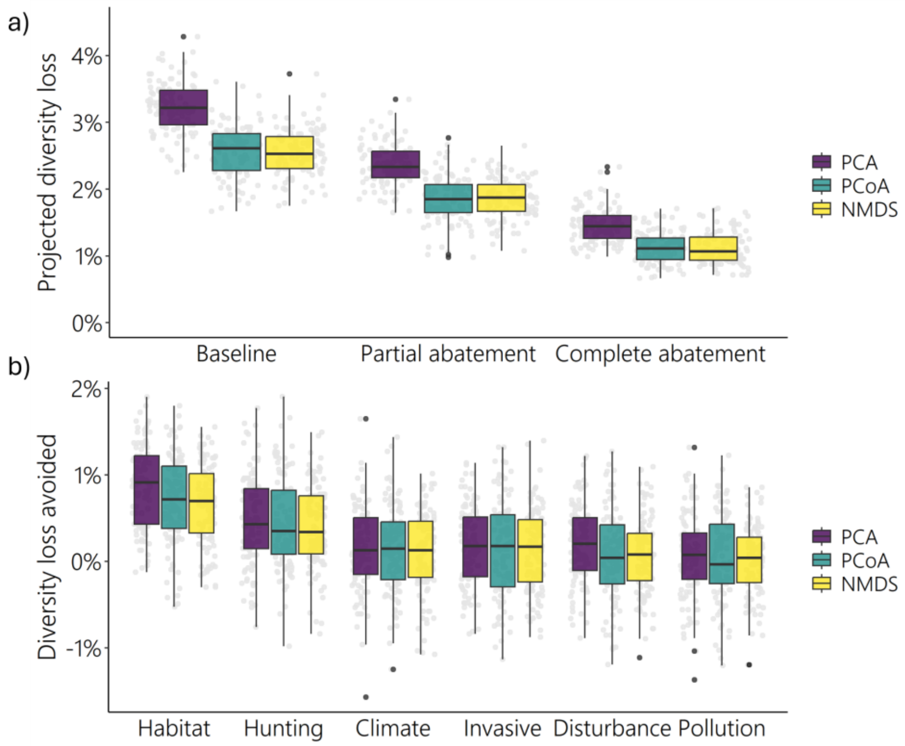


**Supplementary Figure 9. Impact of ordination method on functional richness estimations.** Functional richness was estimated from trait space built using principal component analysis (PCA), principal coordinates analysis (PCoA) and Nonmetric Multidimensional Scaling (NMDS). Grey points show raw data with 100 iterations for each extinction scenario. a) Projected diversity loss under baseline, partial abatement and complete abatement scenarios, and b) Diversity loss avoided under driver-specific abatement. Error bars of diversity loss avoided can go below zero if some iterations of driver-specific abatement have greater projected functional diversity loss than some iterations of baseline extinction scenarios.

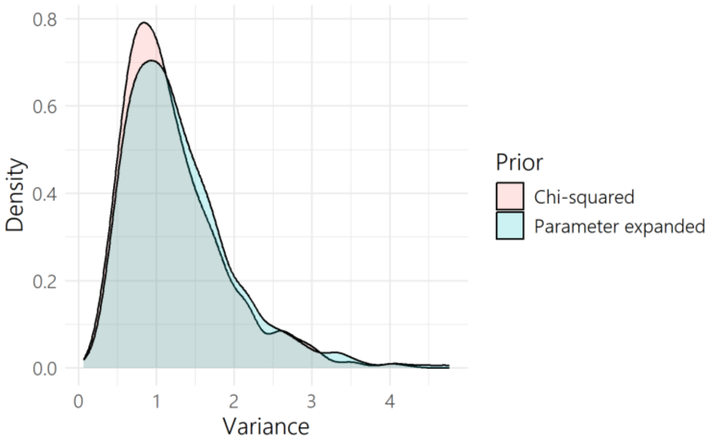


**Supplementary Figure 10. Posterior distribution of phylogenetic variance in the extinction risk model when using a Chi-squared or parameter expanded prior**. The extinction risk model described extinction risk category with threats affecting Near-Threatened and threatened species (2087 species) as fixed effects (Cauchy-scaled priors) and phylogeny and three spatial variables (minimum latitude, maximum latitude and centroid latitude). For the spatial random effects parameter expanded priors were used. MCMC chains were run for 103 000 iterations with a burn-in period of 3000.

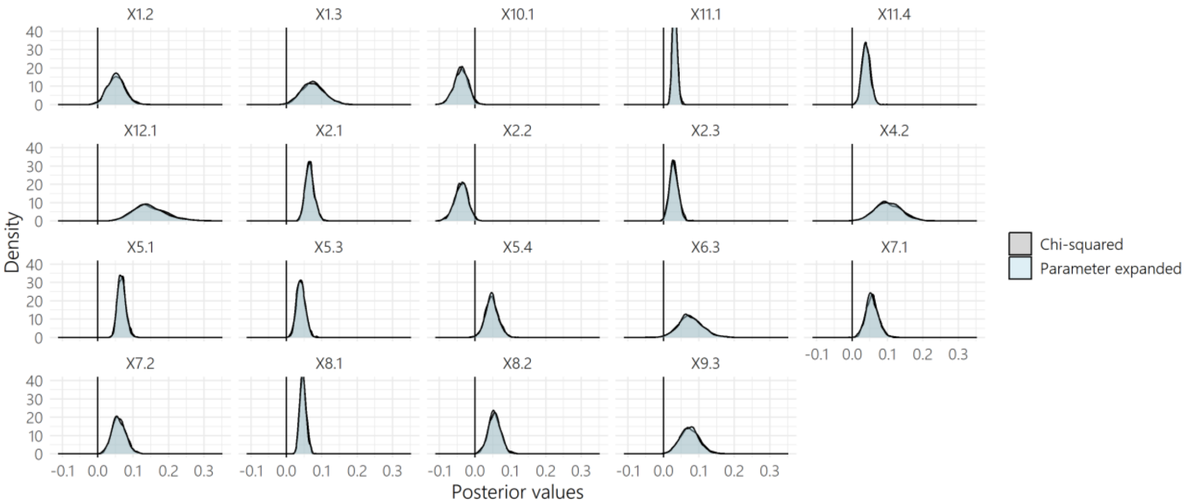


**Supplementary Figure 11. Posterior distributions of fixed effects were similar with different phylogenetic random effect prior specifications.** Posterior estimates (1000) of fixed effects in MCMCglmm model of extinction risk with threats as fixed effects (Cauchy-scaled prior), using an Chi-squared prior (grey) compared to a parameter expanded prior (blue) for the phylogenetic random effect. X1.2 refers to the expected percentage population decline over a 10-year period or three generations as a result of IUCN second order threat "*1.2 Commercial and Industrial Areas*". MCMC chains were run for 103 000 iterations with a burn-in period of 3000.

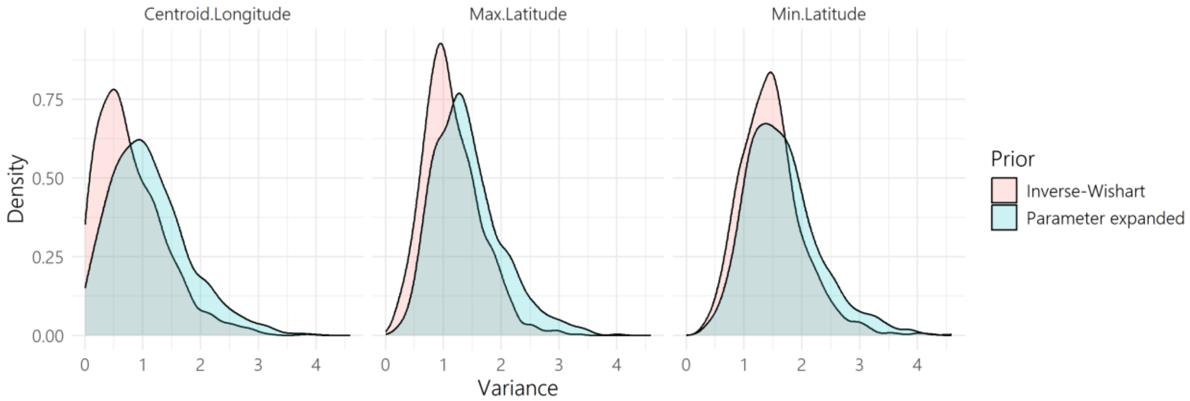


**Supplementary Figure 12. Posterior distribution of spatial variance components in the extinction risk model when using inverse-Wishart or parameter expanded priors.** The extinction risk model described extinction risk category with threats affecting Near-Threatened and threatened species (2087 species) as fixed effects (Cauchy-scaled priors) and phylogeny and three spatial variables (minimum latitude, maximum latitude and centroid latitude). For the phylogenetic random effect a Chi-squared prior was used. MCMC chains were run for 103 000 iterations with a burn-in period of 3000.

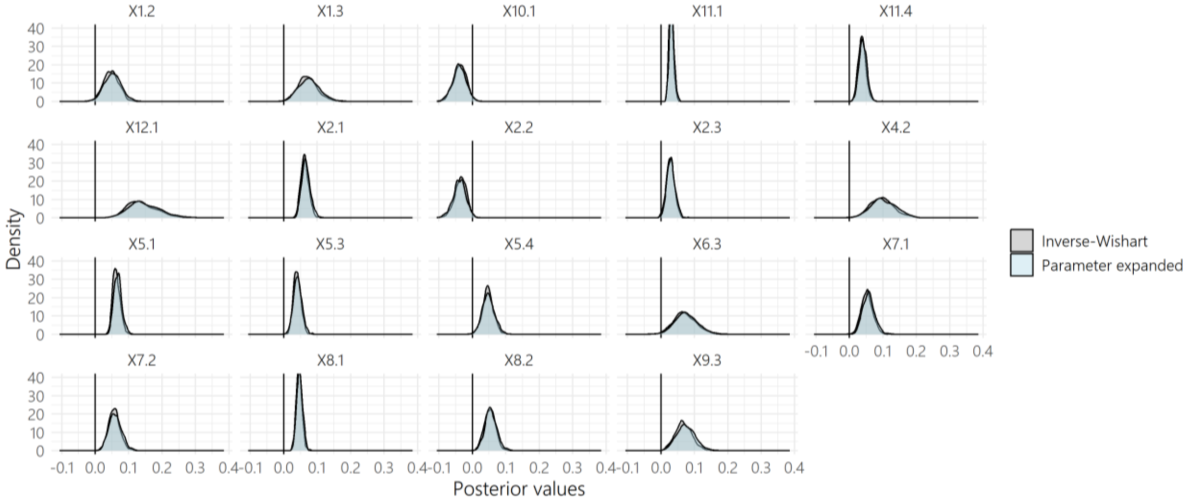


**Supplementary Figure 13. Posterior distributions of fixed effects were similar with different spatial random effect prior specifications.** Posterior estimates (1000) of fixed effects in MCMCglmm model of extinction risk with threats as fixed effects (Cauchy-scaled prior), using an Inverse-Wishart prior (grey) compared to a parameter expanded prior (blue) for spatial random effects (minimum latitude, maximum latitude, centroid longitude). X1.2 refers to the expected percentage population decline over a 10-year period or three generations as a result of IUCN second order threat "*1.2 Commercial and Industrial Areas*". MCMC chains were run for 103 000 iterations with a burn-in period of 3000.

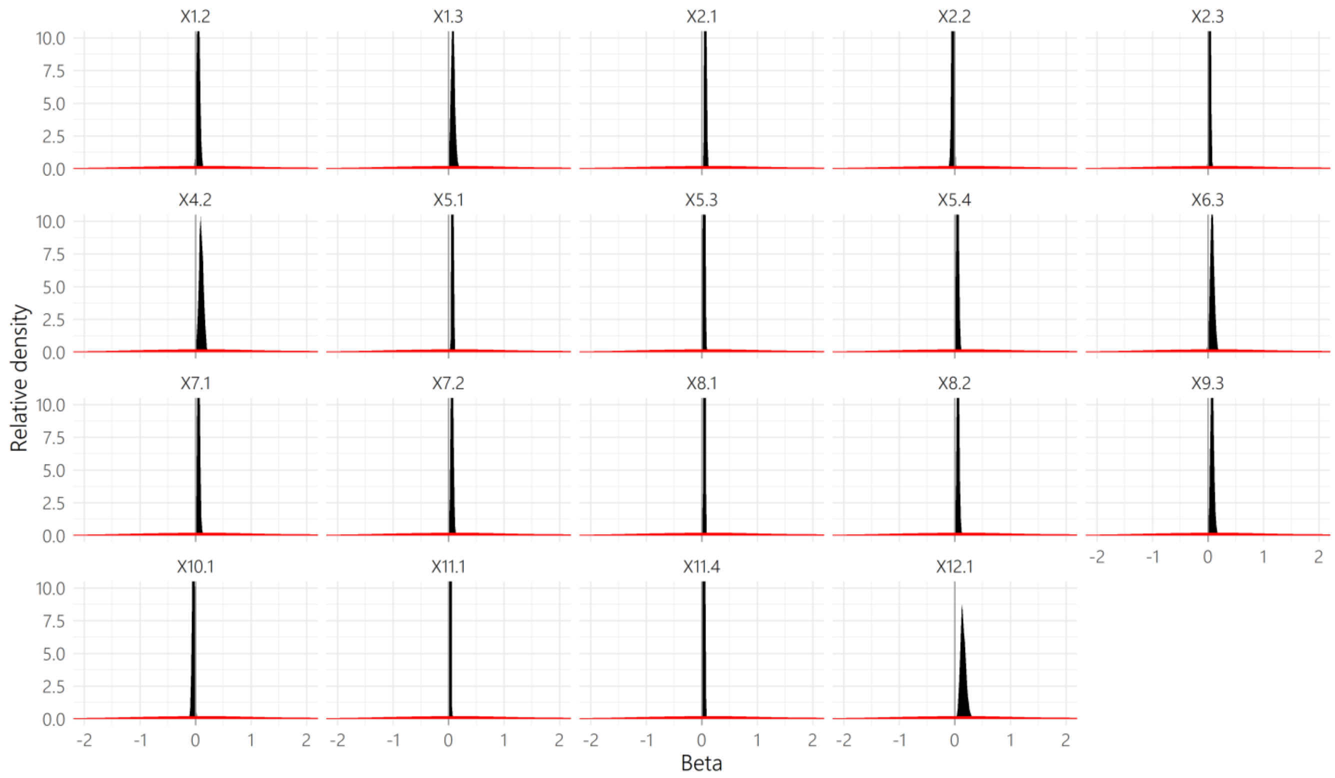


**Supplementary Figure 14. Posterior distributions of fixed effects were not unduly influenced by priors.** Posterior estimates (1000) of fixed effects (black) in MCMCglmm model of extinction risk with threats as fixed effects (Cauchy-scaled prior), where X1.2 refers to the expected percentage population decline over a 10-year period or three generations as a result of IUCN second order threat "*1.2 Commercial and Industrial Areas*". Red shows distribution of Cauchy-scaled priors. Phylogenetic and spatial variables were included as random effects (Chi-squared prior and parameter expanded priors respectively). MCMC chains were run for 103 000 iterations with a burn-in period of 3000.
